# Social environment drives sex and age-specific variation in *Drosophila melanogaster* microbiome composition and predicted function

**DOI:** 10.1101/2020.01.07.895631

**Authors:** Thomas Leech, Laurin McDowall, Kevin P Hopkins, Steven M Sait, Xavier A. Harrison, Amanda Bretman

## Abstract

Social environments influence multiple traits of individuals including immunity, stress and ageing, often in sex-specific ways. The composition of the microbiome (the assemblage of symbiotic microorganisms within a host) is determined by environmental factors and the host’s immune, endocrine and neural systems. The social environment could alter host microbiomes extrinsically by affecting transmission between individuals, likely promoting homogeneity in the microbiome of social partners. Alternatively, intrinsic effects arising from interactions between the microbiome and host physiology (the microbiota-gut-brain axis) could translate social stress into dysbiotic microbiomes, with consequences for host health. We investigated how manipulating social environments during larval and adult life-stages altered the microbiome composition of *Drosophila melanogaster* fruit flies. We used social contexts that particularly alter the development and lifespan of males, predicting that any intrinsic social effects on the microbiome would therefore be sex-specific. The presence of adult males during the larval stage significantly altered the microbiome of pupae of both sexes. In adults, same-sex grouping increased bacterial diversity in both sexes. Importantly, the microbiome community structure of males was more sensitive to social contact at older ages, an effect partially mitigated by housing focal males with young rather than co-aged groups. Functional analyses suggest that these microbiome changes impact ageing and immune responses. This is consistent with the hypothesis that the substantial effects of the social environment on individual health are mediated through intrinsic effects on the microbiome, and provides a model for understanding the mechanistic basis of the microbiota-gut-brain axis.

**Significance statement:** The social environment has pervasive, multifaceted effects on individual health and fitness. If a host’s microbiome is sensitive to the social environment then it could be an important mediator of social effects, as the reciprocal relationships between hosts and their microbiomes have substantial implications for host health. Using a *Drosophila melanogaster* fruit fly model we show that the fly microbiome is sensitive to the social environment in a sex, age and life-stage dependent manner. In particular, older adult male microbiome communities are altered by same-sex social contact, but this depends on the age of the social partners. These changes have functional effects on fly immunity and lifespan, evidence that indeed this is an influential mediator of social effects on health.

## Introduction

Social environments have multiple effects on individual health, including immune responses (1, 2), ageing and ultimately lifespan (3–5). Indeed meta-analyses show that adverse social environments are a health risk factor on a par with obesity and smoking (6). Effects of social environments are complex. They are seen in animals not usually thought of as social, there are marked sex differences, and hence what constitutes a stressful social environment is not straightforward (1, 4, 5). For example, periods of social isolation can be beneficial even in gregarious species (7). The mechanisms that translate information about the social environment into these effects are unclear, but it has been suggested that the microbiome (the community of microorganisms living symbiotically with a host) plays a role (5). Social impacts on microbiomes are expected given that close contact aids horizontal transmission of microbes (8, 9) and social partners will often have similar diets, a key driver of microbiome composition (10). Such extrinsic processes would lead to greater homogeneity in the microbiome of social partners, but would not necessarily have any fitness consequences for the host. However, there is a great deal of interaction between the microbiome and host immune pathways, hormones and neurotransmitters known as the ‘microbiota-gut-brain axis’ (11). Therefore host social environments that impact stress and immune responses (1, 2, 12) could indirectly alter the microbiome. This could have profound consequences for host health given the microbiomes influence on development and behaviour (13), susceptibility to pathogens (14), ageing (15, 16) and fitness trade-offs (17). Therefore, social stress that drives dysbiosis could mediate the effects of social environments on lifespan.

So far the influence of host social interactions on microbiome composition has been investigated exclusively in mammals. Similarities in microbiomes driven by cohabitation, social group membership or social networks seen in ring-tailed lemurs (*Lemur catta*) (18), wild baboons (*Simia hamadryas*) (19) and humans (20) likely represent extrinsic effects of social environments. In mice, social stress alters gut immune gene expression and their gut microbial community (21). Moreover, fecal transfers from mice stressed through isolation recapitulates isolation behaviours in non-isolated mice (22). These studies in mice are suggestive of intrinsic mechanisms connecting host social environments and the microbiome. To broaden our understanding of these effects, we used an invertebrate model system in which simple experimental manipulations of social contact alter ageing and lifespan.

Work in *Drosophila melanogaster* fruit flies has demonstrated multiple effects of social environments on individual behaviour and physiology. We chose to focus on social conditions to which males are particularly sensitive, therefore extrinsic effects of the social environment should affect both sexes equally but intrinsic effects would be seen to a greater extent in males. In adults, same-sex social contact has sex-specific impacts on actuarial and functional senescence (4, 5). Male lifespan is reduced disproportionately by the presence of same-sex cohabitants, especially when given an immune challenge (4), but both sexes can survive longer post-infection with certain bacteria if held with same-sex partners (1). Males use the presence of other males as a cue of potential sperm competition, making sophisticated adjustments to their reproductive behaviour and ejaculate (23, 24). During development, larval density can alter growth rates and adult body size, and the prior presence of adults on food substrates can increase larval survival (25). In addition, when food resources are not limiting, both higher density and the presence of adult males (cues of future sperm competition) stimulate males to develop larger accessory glands (26), and males raised at lower density are better at learning when adult (27). The fly microbiome affects a range of traits including development (28), metabolism (29), immune responses (30) and longevity (16). The fly microbiome is relatively simple (31) and its composition changes across life stages and ages (32). Differences in microbiome community are driven by the environment, for example wild-caught versus laboratory rearing, or maintenance on different food sources (33). Larvae gain gut microbes through ingestion of their egg casing and from their food, and this environmental replenishment continues during adulthood (30), so extrinsic effects of the social environment are likely. Additionally, fly gene expression is socially sensitive, including immune, stress and lifespan related genes (1, 12), so there is potential for intrinsic effects of social environments acting through the microbiota-gut-brain axis.

We captured the bacterial component of the microbiome using 16S sequencing, but for brevity hereafter refer to this as the microbiome. We examined the effect of larval rearing density or presence of adult males, conditions that alter development (25–27), on the microbiome of pupae and one day old adults. As the *D. melanogaster* microbiome is dependent on regular replenishment from ingesting bacteria from the environment, potentially from excreta from other flies (30), we expected that larvae developing in high densities or kept with adults would show greater species richness and changes in microbiome composition. In adults we compared socially isolated flies to those kept in co-aged same sex groups, conditions that alters lifespan in a sex-specific manner (4, 5). In addition, we investigated the effect of the age of the cohabitants by housing an ageing focal fly with a group of consistently young flies, as the effect of social contact on ageing in males can be altered by the age of the partner flies (34). In light of our findings, and because of the importance of microbiomes in combatting infections (30), we tested the ability of adult flies to survive an oral infection.

## Results and discussion

### The presence of adults during development alters the microbiome of pupae

We measured this at the end of development when flies could be sexed, before and after metamorphosis (pupae and 1 day old adults). In pupae, being raised in the presence of adults increased species richness measured as alpha diversity (effect of adult presence F_1, 77_ = 4.648, p = 0.034; effect of life stage F_1, 78_ = 31.39, p <0.001; Figure 1A). There was no effect of sex (Table S1). This is echoed in community structure (beta diversity) where we detected an interaction life stage and adult presence (PERMANOVA F_1, 79_ =7.20, p < 0.001). Distinct separation occurred in the bacterial communities of adult presence and absence groups in the pupal stage (Figure 1B), but not in the 1-day-old adults (Figure 1C). Again, there was no effect of sex (Table S2). *Lactobacillus plantarum, L. brevis* and *Corynebacterium* sp. in particular exhibited differential abundances dependent on life stage and the presence of adults (Table S3).

**Figure 1.**
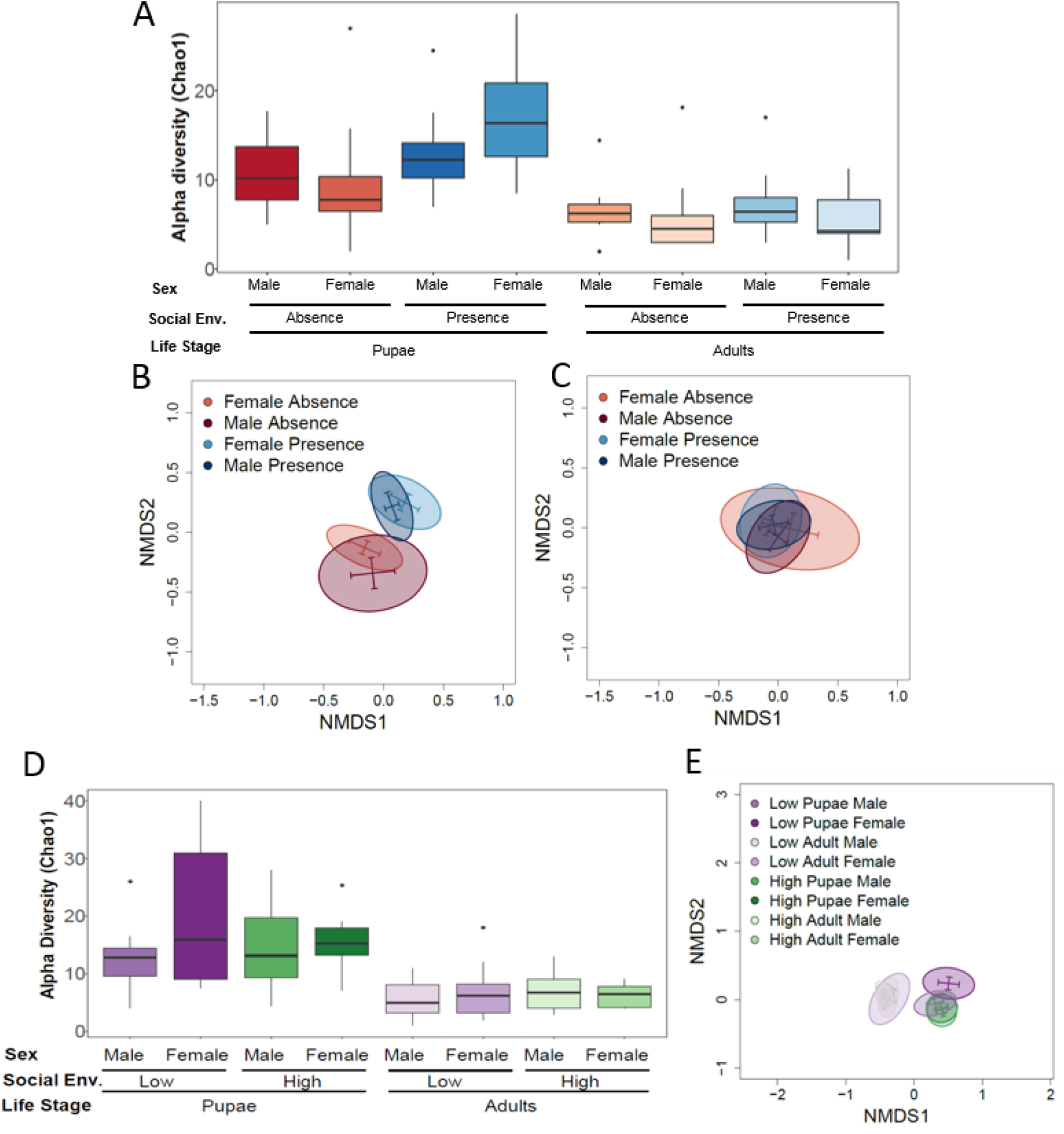
The presence of adults, but not larval density, during larval development alters fly microbiomes. A-C) Larvae were reared in the “Absence” or “Presence” of adult male flies or D-E) were reared low (20) or high (200) density. Flies were sampled as “Pupae” or 1-day-old “Adults”, with males and females analysed separately., Microbiome composition was measured as (A and D) species richness (alpha diversity using the Chao1) and community structure (beta diversity visualised as NMDS plots using Bray-Curtis Dissimilarity Index with 95% confidence ellipses) for pupae (B) and 1 day old adults (C) separately for those raised in the presence or absence of adults, or E) all larval density groups together.

There were no effects of larval density on microbiome composition, but again we observed differences between life stages. Pupae generally displayed a greater species richness (alpha diversity) than their 1-day-old adult counterparts (F_1, 77_ = 35.37, p <0.001; Figure 1D) irrespective of density or sex (Table S4). Likewise, community structure (beta diversity) shows distinction between pupae and 1 day old adult flies (F_1, 78_ = 4.52, p <0.001; Figure 1E), but this was not affected by density or sex (Table S5). Pupae showed increases in *Staphylococcus* sp., *Lactococcus* subsp. *lactis* and *Lactobacillus* sp. compared to adults (Table S6).

Our prediction that more complex social environments would impact microbiome composition was only borne out for the manipulation of adult presence. We chose these social manipulations as they signal future sperm competition to males, hence induce differences in male development and are potentially stressful for males (26, 27). However, their effects on development are not identical (26), (27), suggesting that they convey different social information, and so perhaps it is unsurprising that their effect on the microbiome is likewise not the same. The lack of sex differences in the microbiome at this stage suggests that the underlying mechanism is not associated with the (potentially costly) alterations in development of males to signals of future mating competition (26). We cannot rule out that there was an effect of horizontal transfer from the adults, especially as the presence of adult females improves larval survival partly through inoculating the substrate with yeasts that are an important component of larval diet (25).

Regardless of sex or social manipulation, we found that pupae had a greater species richness than young adults, in line with results observed by Wong et al. (32). This is perhaps unsurprising given that pupae undergo large modifications before eclosion, including expression of antimicrobial peptide genes (35), which may regulate the bacterial community (31), decreasing the number of bacterial taxa observed (32).

### Adult social environment alters microbiome composition

We found that the effect of group housing on the microbiome of adult flies was dependent on age and sex. In 11-day-old flies, bacterial species richness (alpha diversity) was unaffected by social environment and sex (Table S7; Fig 2A). Likewise community structure (beta diversity) was unaffected by social environment, but males and females had distinct communities (Table S8; Fig 2B). However, in 49-day-old flies, bacterial richness was significantly affected by social environment (F_1, 46_ = 8.699, p = 0.0007) with co-aged groups having higher richness compared to single flies or those in mixed-age groups (Table S7; Fig 2C). Community structure was driven by an interaction between social environment and sex (F_1, 47_ = 12.920, p < 0.0001; Fig 2D). To understand this interaction further, we split the data by sex and found that in males there was a highly significant effect of social environment on community structure (F_1, 22_ = 14.054, p < 0.0001), but not in females (F_1, 22_ = 2.188, p = 0.099).

**Figure 2.**
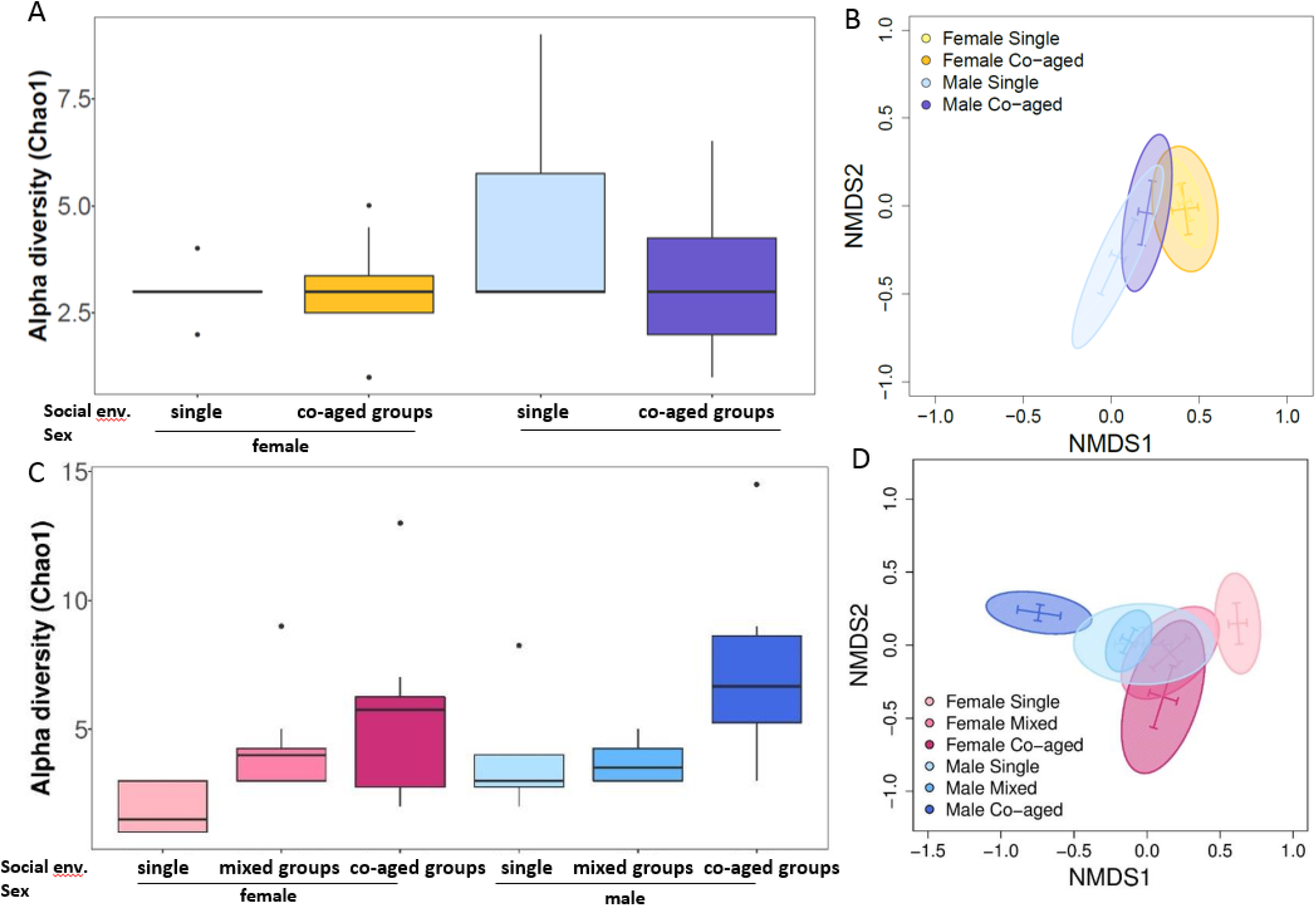
Group housing affects the microbiome of older adult flies. Flies were housed singly or in same-sex groups and were harvested at 11 days (A-B) or 49 days (C-D) post eclosion. For 49 day old flies, groups were either “Co-aged” with the focal fly or were 1-7 days old (“Mixed”). Microbiome composition was measured as (A and C) species richness (alpha diversity using the Chao1) and (B and D) community structure (beta diversity visualised as NMDS plots using Bray-Curtis Dissimilarity Index with 95% confidence ellipses).

There was no significant effect of social environment on relative levels of individual bacterial species, though there were effects of sex and age. Females have significantly lower levels of *Lactobacillus plantarum* and *L. brevis* compared to males (Table S9). Effects of age were only observed in males (Table S10) with young flies having significantly less *L. plantarum* and *L. brevis* than old flies.

These patterns indicate that extrinsic factors, such as shared diet or direct bacterial transfer are unlikely to be solely responsible for the patterns we observe, as these ought to affect males and females equally. Previous work has shown that sex differences in the microbiome become apparent in older adult flies (32) and the effect of the microbiome on fly metabolism is sex-specific (29). The social manipulation we used causes sex differences in lifespan, suggesting that it is more stressful for males than females, or prompts differential investment in physiological processes underlying lifespan-reproduction trade-offs (4, 5). There is increasing evidence for a reciprocal relationship between host stress responses and the microbiome (36), and one direct source of social stress is aggressive interactions. In mice, aggression between males affects colonic mucosa-associated bacterial communities, reducing the relative abundance of key genera including *Lactobacillus* (21). In *D. melanogaster*, males are more aggressive to each other than females, however we have previously been unable to relate levels of aggression to sex-specific patterns in senescence (4, 37). Males respond to sexually competitive environments by increasing mating duration and therefore reproductive fitness (23), but this comes at the cost of lifespan and successful later-life mating attempts (37). If investment in reproduction trades-off with immunosenescence, the result could be quicker ageing and more severe microbial dysbiosis in grouped males. However, neither of these scenarios explain why the effect of grouping on male microbiomes can be ameliorated by housing with young males. There is some evidence that the age of social companions has differential effects on ageing profiles. Males carrying a mutation in the antioxidant enzyme *Sod* have extended lifespan if housed with young males, perhaps because young social partners increased the activity of the focal flies (34). Whether this increased activity drives the extension of lifespan or is a symptom of a less stressful social context, and how this relates to the fly microbiome, remains unclear. However, we are cautious about drawing further conclusions as, due to logistical reasons, our mixed-age treatment were novel to the focal fly whereas the co-aged groups were not. Further tests are required to distinguish fully between the effect of social partner age and social familiarity.

The effects of same-sex social contact on male behaviour, ejaculate and gene expression can be observed on a timescale of hours to a few days (12, 23, 24). However, we observed no effect on the microbiome of young flies, but rather only at older ages, in line with declines in functions such as mating success (37) and climbing ability (4). In *D. melanogaster*, microbial abundance increases with age (16), with all bacterial taxa increasing significantly and resulting in distinct shifts in microbial community structure as the flies age (15). One explanation for the lack of observed differences in young flies may be that the effects of social stress only become apparent as the flies senesce and gene expression becomes less tightly controlled, allowing unchecked proliferation of gut bacteria that impacts gut homeostasis (15, 16). Such a cumulative rather than acute effect of social contact would again be suggestive of intrinsic effects of the social environment acting through the microbiota-gut-brain axis.

### Socially-driven changes in microbiomes likely affect host ageing and immunity

To assess predicted functional implications of changes in the microbial community, we made targeted pair-wise comparisons based on the results of the diversity analysis. These revealed numerous functional pathways that were differentially enriched depending on sex, age and social environment (Tables S11-16). For illustration, we chose five pathways of interest involved in ageing and immunity, which were commonly differentially represented in our data.

In manipulations of larval social environment, the presence of adults had significant effects on the enrichment of these pathways, more so in pupae than in 1-day-old adults, reflecting the findings in terms of microbiome composition, (Fig 3A; Table S11). We found that in pupae, adult presence increased the differential abundances of the *FoxO* and longevity pathways, but decreased abundance of the apoptosis pathway. Further investigation is required to understand the consequences of this, but it is possible that if these alter developmental trajectories (e.g. through *FoxO* activity (39)) they could have long lasting effects even though microbial community alteration itself did not carry-over into adulthood. Indeed, we found that the presence of adults reduced lifespan (Fig 3B, Cox PH *X*^2^_1_ = 6.545, *p* = 0.011) whereas larval density had no effect (Fig 3C, Cox PH *X*^2^_1_ = 1.266, *p* = 0.261), though in both experiments females lived longer than males (Adult presence Cox PH *X*^2^_1_ 109.27, p<0.001; Larval density Cox PH *X*^2^_1_ = 107.56, *p*<0.001). This echoes findings in adult social environments, where treatments showing differences in lifespan (same-sex contact reducing lifespan more in males) are also those showing alterations in their microbial community.

**Figure 3.**
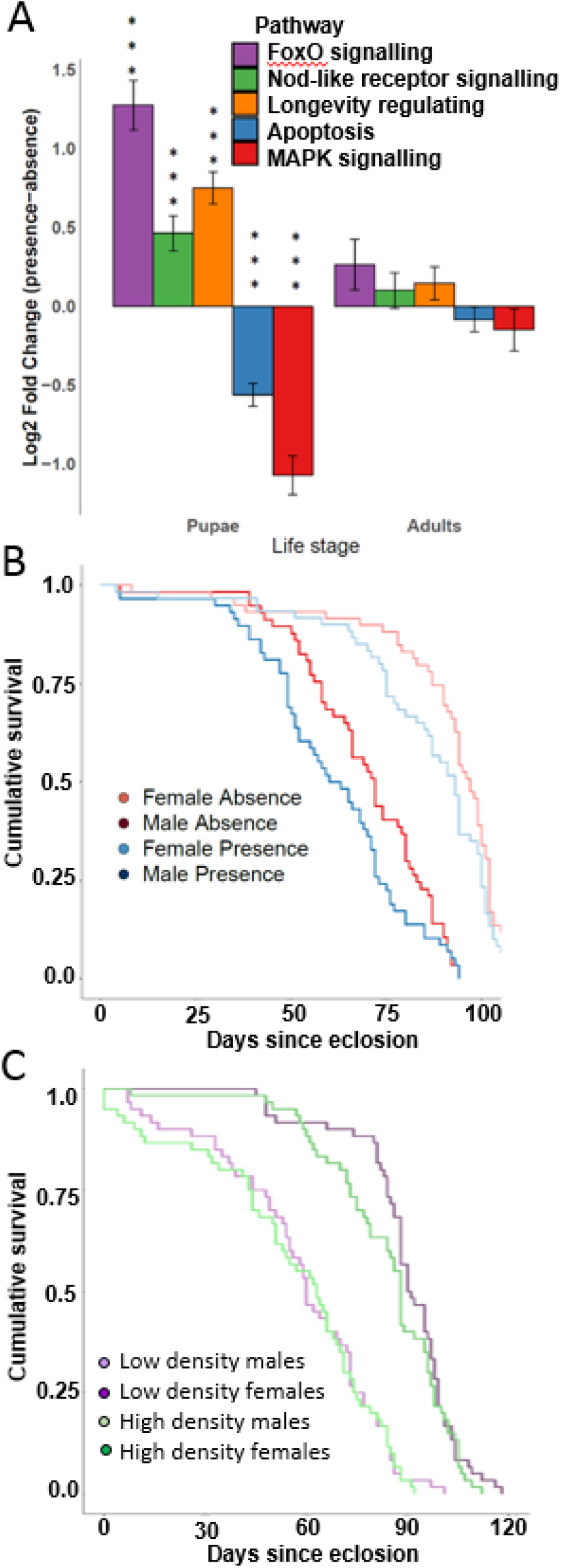
Larval social environment drives microbiome composition changes on functional pathways associated with ageing and alters lifespan. **A)** Predicted microbial effects on gene function was determined using Piphillin, assigned using the KEGG database, and differentially abundant pathways identified by DESeq2 analysis. Comparisons were made for each life stage (pupae or 1 day old adults) between flies reared as larvae with or without adult males present. *** p < 0.001 after Benjamini-Hochberg correction for multiple testing. Lifespan of male and female flies raised (B) in the absence or presence of adults and (C) at low or high density.

In adults, there was a general picture of grouping exacerbating differences in functional pathway abundance between sexes, and for males, differences between young and old flies. Females were largely enriched for these pathways compared to males, but more so in co-aged groups (Figure 4A, Table S12-13). In males, young flies were largely enriched for these pathways compared to old flies, and this was again more prominent in co-aged groups (Figure 4B, Table S14-15). In old males, single flies were more enriched compared to co-aged flies, but not mixed aged groups (Figure 4C, Table S16). This analysis is consistent with our hypothesis that the microbiome mediates the social environmental effect on lifespan and ageing. However, it should be noted that whilst we highlight these as pertaining to our central theme of social effects on lifespan, there were multiple other significantly differentially represented pathways.

**Figure 4.**
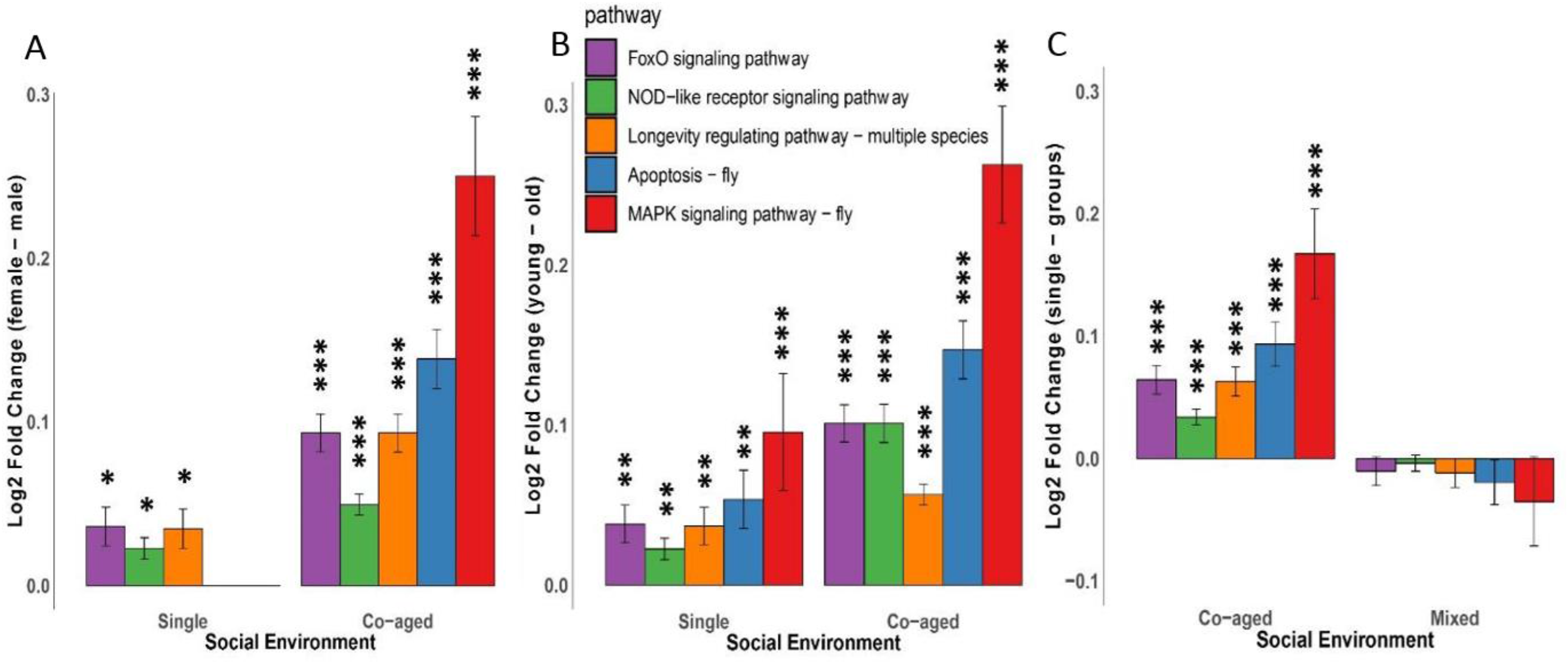
Socially-driven microbiome composition alters host functional pathways associated with ageing. Predicted microbial effects on gene function was determined using Piphillin, assigned using the KEGG database, and differentially abundant pathways identified by DESeq2 analysis. Comparisons were made between (A) 49-day-old females and males that had been housed singly or in co-aged groups (B) 11-day-old and 49 day-old males that had been held singly or in co-aged groups and (C) 49-day-old males held in co-aged groups or with younger flies in mixed age groups. Significant differences * p < 0.05, ** p < 0.01, *** p < 0.001, corrected for multiple testing using the Benjamini-Hochberg method.

Clearly a substantial amount of work is needed to understand the consequences of differential enrichment of these functional pathways. As a starting point we carried out an oral infection assay, as a healthy microbiome, and in particular the presence of *L. plantarum*, can protect against infections (30). We have previously shown that social contact can increase survival after infection (1), however our mode of infection was injection, which therefore bypassed the gut microbiome. We predicted that if social contact caused dysbiosis then we would find post-infection survival reduced if the infection was orally acquired. Indeed we found that isolated males had greater survival after oral infection with *Pseudomonas fluorescens* than grouped males (*X*^2^_1_ = 8.294, *p* = 0.004; Figure S1A), but there was no social effect in females (*X*^2^_1_ = 0.699, *p* = 0.403), mirroring the patterns in the microbial community. However, we could not link this to alterations of particular bacterial species, i.e. differences in abundance of the protective *L. plantarum*. We tested whether this could be driven by males ingesting more of the pathogen. Paired males did not eat more than those held singly so it is unlikely that fewer survived because they consumed more infected food (*X*^2^_1_ = 14.312, *p* = 0.852; Figure S1B). We also found that paired females ate more than single females (*X*^2^_1_ = 25.375, *p* = 0.044), and this social effect on appetite deserves further investigation. In combination with our predicted gene function analysis this indicates that changes in the microbiome could explain why males are susceptible to the immunological and longevity costs of same-sex social contact.

### Conclusions

The social environment has distinct effects on microbiome composition in *D. melanogaster* that are context-dependent. During early life, larval density does not appear to affect microbiome composition, however, the presence of adult males increases diversity in pupae. Whilst this microbial community difference does not carry over into 1 day old adult flies, flies raised in the presence of adults do show shorter lifespans. In adults, same-sex social contact disproportionately affected the microbiome of males, but only in old flies, raising the possibility that immunosenescence is playing a key role. The changes in the microbiome were associated with differential expression of immune and longevity pathways, and male flies housed in same-sex pairs are less able to cope with oral infection, pointing to fitness consequences of these shifts in the microbiome composition. Intriguingly, co-housing ageing males with young social partners ameliorated the changes in microbiome community and functional pathways. Whilst we cannot rule out direct effects such as horizontal bacterial transfer, these results indicate that intrinsic mechanisms such as stress or immune responses could drive the changes in the microbiome, which in turn could explain the differences in lifespan under the social conditions tested. As such this *Drosophila* model could prove key to achieving a mechanistic understanding of the drivers and consequences of the “microbiota-gut-brain” axis.

## Materials and Methods

### Fly stocks and maintenance

*Drosophila melanogaster* wild type (strain Dahomey) were raised on standard sugar-yeast agar medium (40). Flies for all experiments were maintained at a constant 25°C and 50% humidity with 12h:12h light:dark cycle. Experimental larvae were raised at a density of 100 larvae (unless otherwise stated) per 7ml vial supplemented with a live yeast. Upon eclosion, virgin adult flies were sexed under ice anaesthesia and transferred to the relevant social environment.

### Larval social environment

Larval density treatments consisted of 20 (low) or 200 (high) larvae per vial on a concentrated medium to prevent food becoming a limiting factor at high density (26, 27). Adult presence/ absence groups were raised at 100 larvae per vial. The adult presence treatment had 20 adult males added to the vial, removed the day before eclosion. Pupae were collected the day before eclosion, and sexed by the presence of sex combs on male legs. Adults were collected within 8 hours of eclosion, and transferred singly to a vial containing fresh food for approximately 24 hours before freezing at −80°C. Each individual originated from a separate larval vial.

### Adult social environment

Adult males and females were kept alone or in same-sex groups consisting of one focal fly and nine cohabitants. Focal flies were given a small wing-clip so that those in groups could be identified. Focal flies were sacrificed at either 11 days old or 49 days old, ages chosen in line with previous work indicating the senescent effects of social environment become apparent at approximately 49 days old (4). For old flies, to assess the effect of co-ageing within groups, cohabitants were either the same age as the focal fly, or were changed weekly for adults that had eclosed the day before (i.e. constantly aged 1-7 days). Food was changed weekly.

### 16s rRNA sequencing and bioinformatics

For sequencing, each biological replicate was a pool of 8 flies (n = 10 per social environment for larval experiments and n= 8 per social environment for adult experiments). DNA was extracted using the Mobio PowerSoil^®^ DNA Isolation Kit and quality checked using NanoDrop (ND-1000) before being sequenced using paired end 250bp v2 chemistry on an Illumina MiSeq (see SI). Post-sequencing bioinformatics were conducted using mothur (version 38.2) (41) as in (42). Detailed information on library preparation, sequencing and bioinformatic protocols are provided in Supplementary Information. The average library size was ~40k reads per sample after passing quality control.

### Microbiome statistical analysis

All statistical analysis was conducted using R v3.3.2 (R Core Team, 2017) using the phyloseq (43), vegan (44), ggplot2 (45), DESeq2 (46) and lme4 (47) packages. Prior to analysis 18 contaminant Operational Taxanomic Units (OTUs) present in the negative controls were removed (48). One female pupal sample from the larval density treatment was identified as an extreme outlier (Grubb’s test p < 0.05) in number of OTUs (suggestive of contamination) and hence was removed from all subsequent analysis. Sequences were rarefied in order to normalise library sizes. For larval density, the data was rarefied to 20,140 sequences, and for adult presence to 22,718. For adult social environment groups, all were rarefied to 10,840 sequences.

Alpha diversity was estimated using the Chao1 species richness indicator (49). Predictors of alpha diversity were analysed using GLM with social environment, sex and life-stage/age as fixed factors. Models were simplified from the full model using Analysis of Deviance (AOD). We visualised differences in bacterial community structure among samples (beta diversity) using Non-metric multidimensional scaling (NMDS) plots of Bray-Curtis distances. We used PERMANOVA (with 1000 permutations) to examine the effects of social environment, sex and life stage/age on bacterial beta diversity. We used DESeq2 (46) to identify OTUs that differed significantly in relative abundance between groups. Where differentially-abundant OTUs were classified only to genus level, we cross-referenced the sequence in the GreenGenes database (50) using BLAST to identify to species level where possible. Differences in inferred bacterial community function based on predicted gene function was performed using Piphillin (51) and the KEGG reference database (May 2017 release) using a sequence identity cut-off of 97%. We identified differentially abundant functional pathways between treatments using DESeq2 (46).

### Effects of larval social environment on lifespan

To examine adult lifespan, a further 60 flies per treatment group were collected from larval social environments as they eclosed, and kept in single sex groups of 10 on fresh yeast-sugar medium. Each day, the number of mortalities was recorded and then removed. Surviving flies were transferred weekly onto fresh food. Differences in lifespan were analysed using a Cox Proportional Hazards model, with sex and social treatment as factors.

### Effects of adult social environment on survival post-oral infection

Males and females were raised singly or with a same sex partner to 50 days old (food and non-focal flies were changed weekly as above) before being starved for 3 h and then infected with *Pseudomonas fluorescens* via feeding with a bacteria/sucrose/yeast solution (see SI). Pairs were used in this experiment, rather than groups of 10, since previous work had shown a single partner is enough to elicit socially-driven changes in both immune responses (1) and ageing patterns (4). Flies were checked for death every 24 h for one week. We also confirmed that any patterns seen were not driven by a difference in amount of food eaten using a CAFE assay (52) (see SI). Since post-infection lifespan data was limited to one week, a chi squared test was used to determine if the number of flies that died differed by sex and social environment.

## Acknowledgments

Thanks to Zahra Nikakhtari and James Rouse for help with fly work. This work was supported by a University of Leeds 110 Anniversary PhD scholarship to TL, a Boothman, Reynolds and Smithells Scholarship to LM, a ZSL Mission Opportunity Fund grant and Royal Society Research grant (RG130550) to XAH and a University of Leeds Academic fellowship to AB.

## Author Contributions

AB, SMS, LM, TL and XH designed the experiments; LM, TL and AB conducted fly work; sequencing was carried out by KH. LM, TL and XH analysed the data. The manuscript was written by all authors.

## Data accessibility

Sequencing data has been submitted to the NCBI Sequence Read Archive (PRJNA565891, PRJNA565929, PRJNA565132), and all other data will be freely accessible from the Leeds Research Data Repository upon acceptance

