## Supplemental Fig S1 and Tables for "Social environment drives sex and age-specific variation in *Drosophila melanogaster* microbiome composition and predicted function"

**This PDF file includes:**

Supplementary text

Figure S1

Tables S1 to S17

SI References

### Supplementary materials and methods

#### DNA extraction and 16S rRNA sequencing

We PCR amplified the V4 region of the 16S gene in triplicate using 5x HOT FIREPol Blend Master Mix and individually barcoded primer sets as detailed in Kozich et al (2013). PCR conditions entailed a denaturation step of 95°C for 15 minutes, followed by 28 cycles of 95°C for 20s, 50°C for 60s and 72°C for 60s, followed by a final extension step of 72°C for 10 minutes. We also amplified a negative sample containing only lab water, and a mock community of 25 bacterial isolates mixed in equal proportions as a positive control. PCR products were visualised on 2% agarose gel with Gel Red to confirm the presence of clean bands of amplicon, and repeat PCRs conducted for replicates where this was not the case. Sample triplicate PCRs were pooled, and individual sample pools were cleaned using Ampure XP bead cleanup (Beckman Coulter, California, USA). Following bead cleanup, 1μl of sample library was pooled into a preliminary library pool for a titration run. We quantified library concentration in triplicate using Qubit fluorometric assay, and checked the library was of the expected insert size using an Agilent 2200 Tapestation (Agilent Technologies, California, USA). We performed a titration run of the preliminary library on an Illumina MiSeq using 300 cycles of the v2 nano chemistry to quantify the relative number of reads per sample. We then made an equimolar pool of each sample library based on the relative index representation in the titration run. We sequenced this final pooled library using 500 cycles of 250bp PE reads on the v2 chemistry.

#### Bioinformatics

We followed the standard bioinformatic pipeline in the MOTHUR 'MiSeq SOP' (https://www.mothur.org/wiki/MiSeq_SOP). Briefly, contigs were assembled from paired end reads and aligned to the SILVA SEED v123 reference database. We removed homopolymer runs >8, filtered overhangs and identified and removed for potential chimeras using UCLUST.  Sequences were clustered into Operational Taxonomic Units (OTUs) at the 97% threshold. The v123 SILVA database was also used for taxonomic assignment of OTUs. We removed sequences assigned as either Archaea, chloroplasts, or mitochondria. OTU sequences, sequences abundance data and sample metadata were exported from MOTHUR along as a 'biom' object for analysis in the software R.

**Oral infection and CAFE assay**

Social interaction does not alter appetite in *D. melanogaster* (1), but in order to confirm this in our system, the CAFE assay was used (1). Males and females were raised singly (males n = 34, females n = 39) or with a same sex partner (males n = 31, females n = 30) to the age of 50 days. The CAFE assay was conducted to ensure that any differences in post-infection lifespan could not be explained solely by differences across social environment or sex in infectious dose consumed. To ensure that only focal flies from social treatment groups could eat the food, non-focals were kept in transparent Eppendorf tubes with a net lid, inside no-food vials. This ensured they were unable to feed, but that focal flies were still able to detect visual, auditory and olfactory cues from the cohabitant. Flies were allowed to feed for 24 hours before the amount of food eaten was measured and normalised using evaporation controls that contained no flies. To test for differences in the amount of food eaten between groups, one was added to all values in order to make them positive and was subsequently analysed using a GLM with quasi-Poisson errors to account for over dispersion. Individual groups of interest were compared to each other using Mann-Whitney U test, since the data was not normally distributed and were corrected for multiple testing using the Bonferroni method. For oral infection assays, *Pseudomonas fluorescens* (DSMZ 50090) was grown for 48 hours at 25ºc, centrifuged, and the bacterial pellet was re-suspended in a solution of 5% sucrose 5% yeast. To administer the bacteria, 30µl was pipetted onto 7ml agar gel ("no-food" vial) before flies from the relevant groups were added into the no-food vial to feed on the bacteria solution. 30µl of fresh bacterial-sucrose solution was added for 3 days, prior to being transferred onto standard SYA and observed every 24 hours until death for 1 week. Sham controls were included and subsequently removed from the analysis since no flies died.

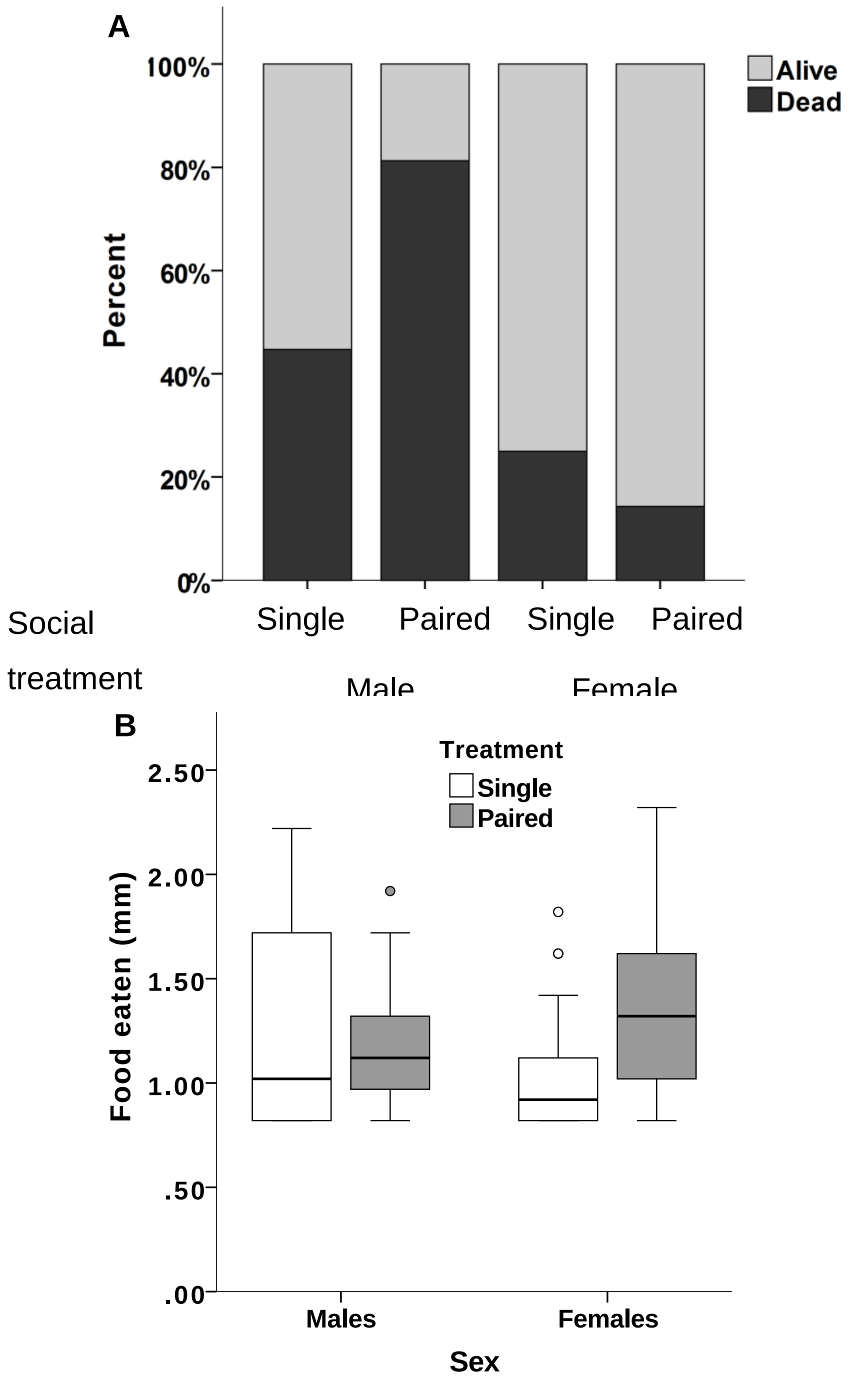


**Figure S1 Effect of same-sex social contact as adults on post-infection survival and appetite.** Flies were kept either alone or in same sex pairs until they were assayed at 52 days old. (A) Percentage of flies that were alive or dead at the end of a one week period**,** after oral infection with *P. fluorescens*. (B) The amount of bacteria-infected food eaten by males and females kept singly and in same-sex pairs.

**Table 1 Effect of adult presence during development on microbiome species richness (alpha diversity).** GLMs were performed using adult presence/absence, life-stage and sex as the explanatory variables for the Chao1 estimate of alpha diversity. Models were simplified from the maximal model using AOD and the values given are the comparison of models with and without the term.

| Explanatory Variable | F | df | *P* value |
|---|---|---|---|
| Adult presence/absence | 4.648 | 1, 77 | 0.0342 |
| Sex | 0.007 | 1, 77 | 0.933 |
| Life-stage | 31.39 | 1, 78 | <0.001 |

**Table 2 Effect of adult presence during development on microbiome community structure.** Beta diversity was analysed using a PERMANOVA with adult presence/absence, life-stage and sex as the explanatory variables.

| Explanatory Variable | F | df | *P* value |
|---|---|---|---|
| Sex | 0.193 | 1, 79 | 0.883 |
| Life-stage*Adult presence/absence | 7.20 | 1, 79 | <0.001 |

**Table 3 Effect of adult presence during development on relative abundance of bacterial species.** Species differential representation was analysed using DESeq2. Life stages were compared within density treatments. Where possible, bacteria were identified to species level using the Greengenes database. P-values are corrected for multiple testing using the Benjamini-Hochberg correction.

| Groups | Bacteria | Log2 Fold Change | Adj. pvalue |
|---|---|---|---|
| Absence Pupae and Adults | *Lactobacillus brevis* | -3.503 | 0.021 |
| | *Lactococcus* subsp. *lactis* | 4.058 | <0.001 |
| | *Lactobacillus* sp. | 3.495 | 0.003 |
| | *Lactobacillus plantarum* | 6.241 | <0.001 |
| Presence Pupae and Adults | *Lactobacillus brevis* | 5.654 | <0.001 |
| | *Lactococcus* subsp. *lactis* | 4.176 | <0.001 |
| | *Lactobacillus* sp. | 3.914 | <0.001 |
| | *Corynebacterium* sp. | 3.655 | <0.001 |
| Presence and Absence Pupae | *Lactobacillus plantarum* | 5.565 | <0.001 |
| | *Lactobacillus brevis* | 10.681 | <0.001 |
| | *Corynebacterium* sp. | 3.386 | <0.001 |

**Table S4 Effect of larval density on microbiome species richness (alpha diversity).** GLMs were performed using density, life-stage and sex as the explanatory variables for the Chao1 estimate of alpha diversity. Models were simplified from the maximal model using AOD and the values given are the comparison of models with and without the term.

| Explanatory term | F | df | *P* value |
|---|---|---|---|
| Density | 0.514 | 1, 76 | 0.475 |
| Sex | 2.696 | 1, 75 | 0.105 |
| Life-stage | 35.37 | 1, 77 | **<0.001** |

**Table S5 Effect of larval density on community structure.** Beta diversity was analysed using a PERMANOVA with density, life-stage and sex as the explanatory variables.

| Explanatory term | F | df | *P* value |
|---|---|---|---|
| Density | 1.277 | 1, 78 | 0.154 |
| Sex | 0.759 | 1, 78 | 0.756 |
| Life-stage | 4.52 | 1, 78 | **<0.001** |

**Table S6 Effect of larval density on relative abundance of bacterial species.** Species differential representation was analysed using DESeq2. Life stages were compared within density treatments. Where possible, bacteria were identified to species level using the Greengenes database. P-values are corrected for multiple testing using the Benjamini-Hochberg correction.

| Groups | Bacteria | Log2 Fold Change | Adj. pvalue |
|---|---|---|---|
| High Density Pupae and High Density Adults | *Staphylococcus* sp. | 3.462 | <0.001 |
| | *Lactobacillus brevis* | -3.567 | <0.001 |
| | *Lactococcus* subsp. *lactis* | 4.773 | <0.001 |
| | *Lactobacillus* sp. | 3.217 | <0.001 |
| Low Density Pupae and Low Density Adults | *Staphylococcus* sp. | 3.574 | <0.001 |
| | *Lactococcus* subsp. *lactis* | 4.244 | <0.001 |
| | *Lactobacillus* sp. | 4.320 | <0.001 |

**Table S7 Effect of same-sex contact in adult flies on microbiome species richness (alpha diversity).** Data were analysed separately for young (11 days post eclosion) and old (49 days post eclosion) flies. GLMs were performed using social environment and sex as the explanatory variables for the Chao1 estimate of alpha diversity. Models were simplified from the maximal model using AOD and the values given are the comparison of models with and without the term.

| Fly age | Explanatory Variable | Test Statistic | df | *P* value |
|---|---|---|---|---|
| Young | Sex | 1.246 | 1, 47 | 0.270 |
| | Social environment | 1.673 | 1, 45 | 0.202 |
| | Social environment*Sex | 1.455 | 1, 45 | 0.234 |
| Old | Social environment | 8.699 | 1, 46 | 0.0007 |
| | Sex | 1.663 | -1, 46 | 0.204 |
| | Social environment*Sex | 1.187 | -1, 45 | 0.315 |

**Table S8 Effect of same sex contact on 11-day old fly microbiome community structure.** Beta diversity was analysed using a PERMANOVA with social environment and sex as the explanatory variables.

| Explanatory Variable | Test Statistic | df | *P* value |
|---|---|---|---|
| Sex | 3.937 | 1, 28 | **0.033** |
| Social environment | 0.349 | 1, 29 | 0.689 |
| Social environment*Sex | 0.508 | 1, 28 | 0.600 |

**Table S9 Effect of sex on relative abundance of bacterial species.** Species differential representation was analysed using DESeq2, comparing males and females of the same age and social environment. Only species exhibiting significant changes in abundance are shown. P-values are corrected for multiple testing using the Benjamini-Hochberg correction.

| social environment | Species | Log2 Fold Change | Adj. *p* value |
|---|---|---|---|
| groups | *Lactobacillus plantarum* | -4.693649 | **0.026** |
| singles | *Lactobacillus plantarum* | -6.436006 | **0.0001** |
| | *Lactobacillus brevis* | -4.067376 | **0.036** |

**Table S10 Effect of age on relative abundance of bacterial species.** Species differential representation was analysed using DESeq2, comparing young and old flies of the same sex and social environment. P-values are corrected for multiple testing using the Benjamini-Hochberg method.

| social environment | Species | Log2 Fold Change | Adj. *p* value |
|---|---|---|---|
| Male groups | *Lactobacillus plantarum* | -6.138006 | **0.0001** |
| | *Lactobacillus brevis* | -5.784371 | **0.0001** |
| Male singles | *Lactobacillus plantarum* | -4.681618 | **0.013** |
| | *Lactobacillus brevis* | -4.395021 | **0.013** |

**Table S11 Effect of adult presence during the larval stage on predicted gene function pathway differential representation from the microbiome of pupae.** P-values are corrected for multiple testing using the Benjamini-Hochberg method.

| KEGG number | Log2 Fold Change | LFC standard error | Adj. *P* value | Pathway Function |
|---|---|---|---|---|
| ko05111 | 3.482 | 0.348 | <0.001 | Biofilm formation - Vibrio cholerae |
| ko02020 | 1.717 | 0.173 | <0.001 | Two-component system |
| ko00300 | -0.646 | 0.068 | <0.001 | Lysine biosynthesis |
| ko00970 | -0.701 | 0.074 | <0.001 | Aminoacyl-tRNA biosynthesis |
| ko03030 | -0.622 | 0.065 | <0.001 | DNA replication |
| ko03440 | -0.691 | 0.073 | <0.001 | Homologous recombination |
| ko03010 | -0.935 | 0.099 | <0.001 | Ribosome |
| ko03430 | -0.531 | 0.057 | <0.001 | Mismatch repair |
| ko00910 | 1.555 | 0.167 | <0.001 | Nitrogen metabolism |
| ko00195 | -1.066 | 0.115 | <0.001 | Photosynthesis |
| ko03018 | -0.565 | 0.061 | <0.001 | RNA degradation |
| ko03020 | -0.773 | 0.084 | <0.001 | RNA polymerase |
| ko00240 | -0.385 | 0.042 | <0.001 | Pyrimidine metabolism |
| ko04066 | -1.421 | 0.155 | <0.001 | HIF-1 signaling pathway |
| ko00550 | -0.541 | 0.059 | <0.001 | Peptidoglycan biosynthesis |
| ko05152 | -0.748 | 0.082 | <0.001 | Tuberculosis |
| ko00710 | -0.747 | 0.082 | <0.001 | Carbon fixation in photosynthetic organisms |
| ko00790 | 1.020 | 0.113 | <0.001 | Folate biosynthesis |
| ko03060 | -0.773 | 0.086 | <0.001 | Protein export |
| ko00785 | -1.008 | 0.113 | <0.001 | Lipoic acid metabolism |
| ko00480 | 0.627 | 0.070 | <0.001 | Glutathione metabolism |
| ko00900 | -0.485 | 0.054 | <0.001 | Terpenoid backbone biosynthesis |
| ko04112 | -0.843 | 0.095 | <0.001 | Cell cycle - Caulobacter |
| ko01120 | 0.509 | 0.058 | <0.001 | Microbial metabolism in diverse environments |

| | | | | |
|---|---|---|---|---|
| ko01051 | -1.075 | 0.122 | <0.001 | Biosynthesis of ansamycins |
| ko00562 | 1.219 | 0.140 | <0.001 | Inositol phosphate metabolism |
| ko00908 | -1.071 | 0.123 | <0.001 | Zeatin biosynthesis |
| ko01055 | -1.071 | 0.123 | <0.001 | Biosynthesis of vancomycin group antibiotics |
| ko04013 | -1.071 | 0.123 | <0.001 | MAPK signaling pathway - fly |
| ko05205 | -1.071 | 0.123 | <0.001 | Proteoglycans in cancer |
| ko03420 | -0.531 | 0.061 | <0.001 | Nucleotide excision repair |
| ko00020 | -0.667 | 0.078 | <0.001 | Citrate cycle (TCA cycle) |
| ko04016 | 1.262 | 0.148 | <0.001 | MAPK signaling pathway - plant |
| ko05014 | 1.534 | 0.181 | <0.001 | Amyotrophic lateral sclerosis (ALS) |
| ko01503 | 1.095 | 0.130 | <0.001 | Cationic antimicrobial peptide (CAMP) resistance |
| ko05010 | -0.719 | 0.086 | <0.001 | Alzheimer disease |
| ko00410 | 2.111 | 0.251 | <0.001 | beta-Alanine metabolism |
| ko00430 | 1.118 | 0.135 | <0.001 | Taurine and hypotaurine metabolism |
| ko00190 | -0.711 | 0.086 | <0.001 | Oxidative phosphorylation |
| ko04260 | -0.955 | 0.116 | <0.001 | Cardiac muscle contraction |
| ko00760 | 1.436 | 0.175 | <0.001 | Nicotinate and nicotinamide metabolism |
| ko00350 | 1.778 | 0.217 | <0.001 | Tyrosine metabolism |
| ko00982 | 1.460 | 0.178 | <0.001 | Drug metabolism - cytochrome P450 |
| ko02026 | 1.900 | 0.233 | <0.001 | Biofilm formation - Escherichia coli |
| ko00521 | 0.792 | 0.098 | <0.001 | Streptomycin biosynthesis |
| ko00380 | 2.174 | 0.270 | <0.001 | Tryptophan metabolism |
| ko04068 | 1.275 | 0.158 | <0.001 | FoxO signaling pathway |
| ko00230 | -0.187 | 0.023 | <0.001 | Purine metabolism |
| ko00980 | 1.301 | 0.163 | <0.001 | Metabolism of xenobiotics by cytochrome P450 |
| ko00280 | 1.111 | 0.140 | <0.001 | Valine, leucine and isoleucine degradation |
| ko00740 | -0.373 | 0.047 | <0.001 | Riboflavin metabolism |
| ko00360 | 2.382 | 0.304 | <0.001 | Phenylalanine metabolism |
| ko04070 | -0.825 | 0.106 | <0.001 | Phosphatidylinositol signaling system |
| ko01212 | 0.534 | 0.069 | <0.001 | Fatty acid metabolism |

| | | | | |
|---|---|---|---|---|
| ko04211 | 1.110 | 0.143 | <0.001 | Longevity regulating pathway |
| ko04214 | -0.560 | 0.073 | <0.001 | Apoptosis - fly |
| ko00650 | 1.070 | 0.139 | <0.001 | Butanoate metabolism |
| ko00564 | -0.282 | 0.037 | <0.001 | Glycerophospholipid metabolism |
| ko04146 | 0.899 | 0.118 | <0.001 | Peroxisome |
| ko01502 | -0.487 | 0.064 | <0.001 | Vancomycin resistance |
| ko05204 | 1.374 | 0.181 | <0.001 | Chemical carcinogenesis |
| ko04932 | -0.702 | 0.093 | <0.001 | Non-alcoholic fatty liver disease (NAFLD) |
| ko05012 | -0.702 | 0.093 | <0.001 | Parkinson disease |
| ko00920 | 2.583 | 0.343 | <0.001 | Sulfur metabolism |
| ko00270 | 0.527 | 0.070 | <0.001 | Cysteine and methionine metabolism |
| ko04072 | -0.857 | 0.115 | <0.001 | Phospholipase D signaling pathway |
| ko05231 | -0.857 | 0.115 | <0.001 | Choline metabolism in cancer |
| ko04626 | 0.835 | 0.112 | <0.001 | Plant-pathogen interaction |
| ko00520 | 0.908 | 0.122 | <0.001 | Amino sugar and nucleotide sugar metabolism |
| ko04213 | 0.752 | 0.102 | <0.001 | Longevity regulating pathway - multiple species |
| ko01524 | 0.810 | 0.110 | <0.001 | Platinum drug resistance |
| ko00330 | 1.676 | 0.229 | <0.001 | Arginine and proline metabolism |
| ko00540 | 5.114 | 0.699 | <0.001 | Lipopolysaccharide biosynthesis |
| ko00903 | 5.189 | 0.714 | <0.001 | Limonene and pinene degradation |
| ko00981 | 5.265 | 0.725 | <0.001 | Insect hormone biosynthesis |
| ko04920 | 6.628 | 0.912 | <0.001 | Adipocytokine signaling pathway |
| ko05016 | -0.568 | 0.078 | <0.001 | Huntington disease |
| ko00404 | 6.247 | 0.870 | <0.001 | Staurosporine biosynthesis |
| ko00565 | 6.173 | 0.863 | <0.001 | Ether lipid metabolism |
| ko01501 | 0.968 | 0.136 | <0.001 | beta-Lactam resistance |
| ko04614 | 5.678 | 0.801 | <0.001 | Renin-angiotensin system |
| ko00053 | 0.888 | 0.126 | <0.001 | Ascorbate and aldarate metabolism |
| ko00770 | 0.922 | 0.131 | <0.001 | Pantothenate and CoA biosynthesis |
| ko00791 | 5.949 | 0.850 | <0.001 | Atrazine degradation |

| | | | | |
|---|---|---|---|---|
| ko04919 | 5.692 | 0.819 | <0.001 | Thyroid hormone signaling pathway |
| ko01040 | 1.089 | 0.157 | <0.001 | Biosynthesis of unsaturated fatty acids |
| ko00640 | 0.514 | 0.074 | <0.001 | Propanoate metabolism |
| ko00401 | 0.797 | 0.116 | <0.001 | Novobiocin biosynthesis |
| ko02010 | 1.125 | 0.164 | <0.001 | ABC transporters |
| ko03320 | 4.587 | 0.668 | <0.001 | PPAR signaling pathway |
| ko05418 | 0.433 | 0.063 | <0.001 | Fluid shear stress and atherosclerosis |
| ko02025 | 5.223 | 0.765 | <0.001 | Biofilm formation - Pseudomonas aeruginosa |
| ko00591 | 5.243 | 0.772 | <0.001 | Linoleic acid metabolism |
| ko05020 | 5.243 | 0.772 | <0.001 | Prion diseases |
| ko00281 | 5.068 | 0.748 | <0.001 | Geraniol degradation |
| ko05206 | 4.929 | 0.739 | <0.001 | MicroRNAs in cancer |
| ko00364 | 5.646 | 0.861 | <0.001 | Fluorobenzoate degradation |
| ko01062 | 4.843 | 0.739 | <0.001 | Biosynthesis of terpenoids and steroids |
| ko00906 | 4.842 | 0.741 | <0.001 | Carotenoid biosynthesis |
| ko00930 | 4.544 | 0.697 | <0.001 | Caprolactam degradation |
| ko04974 | 4.841 | 0.743 | <0.001 | Protein digestion and absorption |
| ko00460 | 0.657 | 0.101 | <0.001 | Cyanoamino acid metabolism |
| ko00130 | -0.434 | 0.067 | <0.001 | Ubiquinone and other terpenoid-quinone biosynthesis |
| ko04122 | 0.442 | 0.068 | <0.001 | Sulfur relay system |
| ko00630 | 0.682 | 0.106 | <0.001 | Glyoxylate and dicarboxylate metabolism |
| ko04011 | 5.399 | 0.839 | <0.001 | MAPK signaling pathway - yeast |
| ko05132 | 5.604 | 0.871 | <0.001 | Salmonella infection |
| ko00600 | 4.259 | 0.665 | <0.001 | Sphingolipid metabolism |
| ko00620 | 0.560 | 0.088 | <0.001 | Pyruvate metabolism |
| ko00623 | 5.522 | 0.866 | <0.001 | Toluene degradation |
| ko01100 | 0.161 | 0.025 | <0.001 | Metabolic pathways |
| ko00440 | 3.477 | 0.548 | <0.001 | Phosphonate and phosphinate metabolism |
| ko04216 | 4.589 | 0.726 | <0.001 | Ferroptosis |

| | | | | |
|---|---|---|---|---|
| ko00311 | 4.573 | 0.734 | <0.001 | Penicillin and cephalosporin biosynthesis |
| ko00642 | 4.812 | 0.773 | <0.001 | Ethylbenzene degradation |
| ko00624 | 4.475 | 0.721 | <0.001 | Polycyclic aromatic hydrocarbon degradation |
| ko04726 | 4.608 | 0.743 | <0.001 | Serotonergic synapse |
| ko04728 | 4.608 | 0.743 | <0.001 | Dopaminergic synapse |
| ko05030 | 4.608 | 0.743 | <0.001 | Cocaine addiction |
| ko05031 | 4.608 | 0.743 | <0.001 | Amphetamine addiction |
| ko05034 | 4.608 | 0.743 | <0.001 | Alcoholism |
| ko00071 | 4.230 | 0.682 | <0.001 | Fatty acid degradation |
| ko00310 | 1.028 | 0.166 | <0.001 | Lysine degradation |
| ko04978 | 4.343 | 0.702 | <0.001 | Mineral absorption |
| ko05219 | 4.344 | 0.702 | <0.001 | Bladder cancer |
| ko02030 | 5.199 | 0.842 | <0.001 | Bacterial chemotaxis |
| ko04113 | 4.345 | 0.705 | <0.001 | Meiosis - yeast |
| ko05133 | 1.000 | 0.163 | <0.001 | Pertussis |
| ko00590 | 4.329 | 0.712 | <0.001 | Arachidonic acid metabolism |
| ko04922 | -0.692 | 0.114 | <0.001 | Glucagon signaling pathway |
| ko00400 | 1.763 | 0.292 | <0.001 | Phenylalanine, tyrosine and tryptophan biosynthesis |
| ko00361 | 3.572 | 0.602 | <0.001 | Chlorocyclohexane and chlorobenzene degradation |
| ko00622 | 3.682 | 0.627 | <0.001 | Xylene degradation |
| ko01053 | 3.432 | 0.585 | <0.001 | Biosynthesis of siderophore group nonribosomal peptides |
| ko00643 | 3.455 | 0.596 | <0.001 | Styrene degradation |
| ko04612 | -1.063 | 0.184 | <0.001 | Antigen processing and presentation |
| ko04657 | -1.063 | 0.184 | <0.001 | IL-17 signaling pathway |
| ko04659 | -1.063 | 0.184 | <0.001 | Th17 cell differentiation |
| ko04914 | -1.063 | 0.184 | <0.001 | Progesterone-mediated oocyte maturation |
| ko04915 | -1.063 | 0.184 | <0.001 | Estrogen signaling pathway |
| ko05215 | -1.063 | 0.184 | <0.001 | Prostate cancer |

| | | | | |
|---|---|---|---|---|
| ko00680 | 0.568 | 0.099 | <0.001 | Methane metabolism |
| ko00592 | 4.482 | 0.787 | <0.001 | alpha-Linolenic acid metabolism |
| ko04964 | 4.482 | 0.787 | <0.001 | Proximal tubule bicarbonate reclamation |
| ko02040 | 5.773 | 1.022 | <0.001 | Flagellar assembly |
| ko00450 | 0.235 | 0.042 | <0.001 | Selenocompound metabolism |
| ko05200 | -0.441 | 0.078 | <0.001 | Pathways in cancer |
| ko00983 | 0.447 | 0.079 | <0.001 | Drug metabolism - other enzymes |
| ko00621 | 3.377 | 0.601 | <0.001 | Dioxin degradation |
| ko01523 | -0.327 | 0.060 | <0.001 | Antifolate resistance |
| ko00524 | 3.024 | 0.559 | <0.001 | Neomycin, kanamycin and gentamicin biosynthesis |
| ko00040 | -0.311 | 0.058 | <0.001 | Pentose and glucuronate interconversions |
| ko00627 | 3.551 | 0.664 | <0.001 | Aminobenzoate degradation |
| ko04727 | 0.518 | 0.098 | <0.001 | GABAergic synapse |
| ko00523 | 0.452 | 0.085 | <0.001 | Polyketide sugar unit biosynthesis |
| ko00660 | 0.643 | 0.122 | <0.001 | C5-Branched dibasic acid metabolism |
| ko05230 | -0.413 | 0.079 | <0.001 | Central carbon metabolism in cancer |
| ko02024 | 0.876 | 0.170 | <0.001 | Quorum sensing |
| ko05120 | -0.492 | 0.096 | <0.001 | Epithelial cell signaling in Helicobacter pylori infection |
| ko05340 | 2.976 | 0.584 | <0.001 | Primary immunodeficiency |
| ko00966 | 2.858 | 0.561 | <0.001 | Glucosinolate biosynthesis |
| ko00362 | 3.665 | 0.722 | <0.001 | Benzoate degradation |
| ko00960 | 0.632 | 0.125 | <0.001 | Tropane, piperidine and pyridine alkaloid biosynthesis |
| ko01230 | 0.345 | 0.070 | <0.001 | Biosynthesis of amino acids |
| ko04930 | 2.583 | 0.528 | <0.001 | Type II diabetes mellitus |
| ko05203 | 2.583 | 0.528 | <0.001 | Viral carcinogenesis |
| ko04916 | 3.429 | 0.706 | <0.001 | Melanogenesis |
| ko00030 | 0.313 | 0.065 | <0.001 | Pentose phosphate pathway |
| ko00525 | -0.339 | 0.071 | <0.001 | Acarbose and validamycin biosynthesis |

| | | | | |
|---|---|---|---|---|
| ko00965 | 3.423 | 0.720 | <0.001 | Betalain biosynthesis |
| ko03450 | 3.423 | 0.720 | <0.001 | Non-homologous end-joining |
| ko05142 | 3.420 | 0.726 | <0.001 | Chagas disease (American trypanosomiasis) |
| ko00072 | 3.012 | 0.640 | <0.001 | Synthesis and degradation of ketone bodies |
| ko05143 | 3.419 | 0.730 | <0.001 | African trypanosomiasis |
| ko04918 | 2.683 | 0.581 | <0.001 | Thyroid hormone synthesis |
| ko01220 | 3.123 | 0.682 | <0.001 | Degradation of aromatic compounds |
| ko01130 | 0.127 | 0.029 | <0.001 | Biosynthesis of antibiotics |
| ko04940 | -0.308 | 0.070 | <0.001 | Type I diabetes mellitus |
| ko03013 | 3.042 | 0.700 | <0.001 | RNA transport |
| ko04910 | 2.497 | 0.580 | <0.001 | Insulin signaling pathway |
| ko04141 | -0.738 | 0.174 | <0.001 | Protein processing in endoplasmic reticulum |
| ko01200 | 0.173 | 0.041 | <0.001 | Carbon metabolism |
| ko04621 | 0.463 | 0.111 | <0.001 | NOD-like receptor signaling pathway |
| ko00260 | 0.251 | 0.060 | <0.001 | Glycine, serine and threonine metabolism |
| ko05322 | 3.195 | 0.773 | <0.001 | Systemic lupus erythematosus |
| ko00625 | 2.676 | 0.648 | <0.001 | Chloroalkane and chloroalkene degradation |
| ko00332 | 2.308 | 0.559 | <0.001 | Carbapenem biosynthesis |
| ko00405 | 2.308 | 0.560 | <0.001 | Phenazine biosynthesis |
| ko04080 | -5.320 | 1.302 | <0.001 | Neuroactive ligand-receptor interaction |
| ko05166 | -5.320 | 1.302 | <0.001 | Human T-cell leukemia virus 1 infection |
| ko00140 | -4.532 | 1.116 | <0.001 | Steroid hormone biosynthesis |
| ko00511 | 2.743 | 0.676 | <0.001 | Other glycan degradation |
| ko00340 | 2.938 | 0.734 | <0.001 | Histidine metabolism |
| ko00513 | -4.399 | 1.105 | <0.001 | Various types of N-glycan biosynthesis |
| ko00604 | -4.399 | 1.105 | <0.001 | Glycosphingolipid biosynthesis - ganglio series |
| ko04622 | -4.399 | 1.105 | <0.001 | RIG-I-like receptor signaling pathway |
| ko04142 | -4.397 | 1.108 | <0.001 | Lysosome |
| ko04210 | 0.321 | 0.081 | <0.001 | Apoptosis |

| | | | | |
|---|---|---|---|---|
| ko00603 | -4.397 | 1.110 | <0.001 | Glycosphingolipid biosynthesis - globo and isoglobo series |
| ko00290 | 2.789 | 0.720 | <0.001 | Valine, leucine and isoleucine biosynthesis |
| ko04917 | 2.354 | 0.633 | <0.001 | Prolactin signaling pathway |
| ko00830 | 2.127 | 0.579 | <0.001 | Retinol metabolism |
| ko00121 | -3.644 | 1.036 | <0.001 | Secondary bile acid biosynthesis |
| ko00860 | -0.216 | 0.063 | <0.001 | Porphyrin and chlorophyll metabolism |
| ko04212 | 0.172 | 0.053 | 0.001637 | Longevity regulating pathway - worm |
| ko00052 | 2.317 | 0.725 | 0.001787 | Galactose metabolism |
| ko00950 | 0.295 | 0.093 | 0.001863 | Isoquinoline alkaloid biosynthesis |
| ko00670 | -0.196 | 0.065 | 0.003221 | One carbon pool by folate |
| ko00531 | 1.665 | 0.572 | 0.004549 | Glycosaminoglycan degradation |
| ko01054 | 1.566 | 0.544 | 0.005019 | Nonribosomal peptide structures |
| ko03008 | -0.270 | 0.095 | 0.005601 | Ribosome biogenesis in eukaryotes |
| ko04152 | 0.302 | 0.113 | 0.009595 | AMPK signaling pathway |
| ko01210 | 0.115 | 0.044 | 0.010516 | 2-Oxocarboxylic acid metabolism |
| ko00626 | 1.609 | 0.616 | 0.011058 | Naphthalene degradation |
| ko03410 | -0.094 | 0.037 | 0.013821 | Base excision repair |
| ko04724 | 0.252 | 0.101 | 0.015489 | Glutamatergic synapse |
| ko00780 | 0.147 | 0.064 | 0.026537 | Biotin metabolism |
| ko00720 | -0.093 | 0.041 | 0.029214 | Carbon fixation pathways in prokaryotes |
| ko05211 | -0.213 | 0.095 | 0.029591 | Renal cell carcinoma |
| ko00471 | -0.095 | 0.045 | 0.042108 | D-Glutamine and D-glutamate metabolism |
| ko04115 | -0.138 | 0.066 | 0.043473 | p53 signaling pathway |
| ko04215 | -0.138 | 0.066 | 0.043473 | Apoptosis - multiple species |
| ko05145 | -0.138 | 0.066 | 0.043473 | Toxoplasmosis |
| ko05150 | -1.885 | 0.903 | 0.043473 | Staphylococcus aureus infection |
| ko05161 | -0.138 | 0.066 | 0.043473 | Hepatitis B |
| ko05164 | -0.138 | 0.066 | 0.043473 | Influenza A |
| ko05168 | -0.138 | 0.066 | 0.043473 | Herpes simplex virus 1 infection |

| | | | | |
|---|---|---|---|---|
| ko05210 | -0.138 | 0.066 | 0.043473 | Colorectal cancer |
| ko05222 | -0.138 | 0.066 | 0.043473 | Small cell lung cancer |
| ko05416 | -0.138 | 0.066 | 0.043473 | Viral myocarditis |

**Table S12 Effect of sex on functional pathway abundance for single adult flies.** Predicted gene function was performed using Piphillin and the KEGG reference database (May 2017 release) using a sequence identity cut-off of 97%. DESeq2 analysis was performed to identify pathways that were differentially abundant between 49-day old single males compared to females. P-values are corrected for multiple testing using the Benjamini-Hochberg method.

| KEGG number | Log2 Fold Change | LFC standard error | Adj. *P* value | Pathway Function |
|---|---|---|---|---|
| ko00906 | -8.347 | 1.425 | <0.0001 | Carotenoid biosynthesis |
| ko04910 | -7.968 | 1.402 | <0.0001 | Insulin signaling pathway |
| ko01062 | -8.331 | 1.510 | <0.0001 | Biosynthesis of terpenoids and steroids |
| ko04622 | -8.409 | 1.569 | <0.0001 | RIG-I-like receptor signaling pathway |
| ko00332 | -5.164 | 1.054 | <0.0001 | Carbapenem biosynthesis |
| ko00405 | -5.232 | 1.069 | <0.0001 | Phenazine biosynthesis |
| ko02025 | -4.874 | 1.058 | <0.0001 | Biofilm formation - Pseudomonas aeruginosa |
| ko00340 | -4.500 | 1.042 | <0.0001 | Histidine metabolism |
| ko00400 | -0.491 | 0.133 | 0.006 | Phenylalanineand tryptophan biosynthesis |
| ko00471 | 0.010 | 0.003 | 0.012 | D-Glutamine and D-glutamate metabolism |
| ko01523 | 0.018 | 0.005 | 0.012 | Antifolate resistance |
| ko03018 | 0.015 | 0.004 | 0.012 | RNA degradation |
| ko04621 | 0.023 | 0.007 | 0.012 | NOD-like receptor signaling pathway |
| ko00240 | 0.020 | 0.006 | 0.015 | Pyrimidine metabolism |
| ko00900 | 0.012 | 0.004 | 0.015 | Terpenoid backbone biosynthesis |
| ko00970 | 0.019 | 0.006 | 0.015 | Aminoacyl-tRNA biosynthesis |
| ko00908 | 0.036 | 0.012 | 0.020 | Zeatin biosynthesis |
| ko03008 | 0.036 | 0.012 | 0.020 | Ribosome biogenesis in eukaryotes |
| ko04724 | 0.036 | 0.012 | 0.020 | Glutamatergic synapse |

| | | | | |
|---|---|---|---|---|
| ko05205 | 0.036 | 0.012 | 0.020 | Proteoglycans in cancer |
| ko04068 | 0.036 | 0.012 | 0.020 | FoxO signaling pathway |
| ko04211 | 0.036 | 0.012 | 0.020 | Longevity regulating pathway |
| ko05014 | 0.036 | 0.012 | 0.020 | Amyotrophic lateral sclerosis (ALS) |
| ko03430 | 0.019 | 0.006 | 0.020 | Mismatch repair |
| ko02026 | -0.185 | 0.061 | 0.021 | Biofilm formation - Escherichia coli |
| ko00051 | -0.228 | 0.077 | 0.024 | Fructose and mannose metabolism |
| ko00920 | -0.170 | 0.057 | 0.025 | Sulfur metabolism |
| ko00280 | 0.032 | 0.011 | 0.025 | Valine, leucine and isoleucine degradation |
| ko00660 | 0.035 | 0.012 | 0.025 | C5-Branched dibasic acid metabolism |
| ko00780 | 0.005 | 0.002 | 0.025 | Biotin metabolism |
| ko04626 | 0.035 | 0.012 | 0.025 | Plant-pathogen interaction |
| ko04066 | 0.027 | 0.009 | 0.026 | HIF-1 signaling pathway |
| ko04213 | 0.035 | 0.012 | 0.026 | Longevity regulating pathway - multiple species |
| ko02060 | -3.063 | 1.099 | 0.033 | Phosphotransferase system (PTS) |
| ko03420 | 0.014 | 0.005 | 0.033 | Nucleotide excision repair |
| ko04141 | 0.024 | 0.009 | 0.033 | Protein processing in endoplasmic reticulum |
| ko00670 | 0.021 | 0.008 | 0.034 | One carbon pool by folate |
| ko00785 | 0.029 | 0.011 | 0.034 | Lipoic acid metabolism |
| ko03010 | 0.023 | 0.009 | 0.037 | Ribosome |
| ko04212 | 0.029 | 0.011 | 0.038 | Longevity regulating pathway - worm |
| ko03030 | 0.024 | 0.009 | 0.042 | DNA replication |
| ko00270 | -0.063 | 0.024 | 0.045 | Cysteine and methionine metabolism |
| ko00564 | 0.022 | 0.008 | 0.047 | Glycerophospholipid metabolism |
| ko05152 | 0.032 | 0.012 | 0.047 | Tuberculosis |

**Table S13 Effect of sex on functional pathway abundance for co-aged grouped adult flies.** Predicted gene function was performed using Piphillin and the KEGG reference database (May 2017 release) using a sequence identity cut-off of 97%. DESeq2 analysis was performed to identify pathways that were differentially abundant between 49-day old males compared to females held in co-aged groups. P-values are corrected for multiple testing using the Benjamini-Hochberg method.

| KEGG number | Log2 Fold Change | LFC standard error | Adj. *P* Value | Pathway Function |
|---|---|---|---|---|
| ko00980 | -0.125 | 0.014 | <0.0001 | Metabolism of xenobiotics by cytochrome P450 |
| ko00982 | -0.125 | 0.014 | <0.0001 | Drug metabolism - cytochrome P450 |
| ko01040 | -0.125 | 0.014 | <0.0001 | Biosynthesis of unsaturated fatty acids |
| ko00908 | 0.093 | 0.011 | <0.0001 | Zeatin biosynthesis |
| ko03008 | 0.093 | 0.011 | <0.0001 | Ribosome biogenesis in eukaryotes |
| ko04724 | 0.093 | 0.011 | <0.0001 | Glutamatergic synapse |
| ko05205 | 0.093 | 0.011 | <0.0001 | Proteoglycans in cancer |
| ko04068 | 0.093 | 0.011 | <0.0001 | FoxO signaling pathway |
| ko04211 | 0.093 | 0.011 | <0.0001 | Longevity regulating pathway |
| ko05014 | 0.093 | 0.011 | <0.0001 | Amyotrophic lateral sclerosis (ALS) |
| ko00910 | -0.163 | 0.020 | <0.0001 | Nitrogen metabolism |
| ko00660 | 0.093 | 0.012 | <0.0001 | C5-Branched dibasic acid metabolism |
| ko04626 | 0.093 | 0.012 | <0.0001 | Plant-pathogen interaction |
| ko00051 | -0.608 | 0.077 | <0.0001 | Fructose and mannose metabolism |
| ko04213 | 0.093 | 0.012 | <0.0001 | Longevity regulating pathway - multiple species |
| ko00480 | -0.034 | 0.004 | <0.0001 | Glutathione metabolism |
| ko05111 | -0.525 | 0.067 | <0.0001 | Biofilm formation - Vibrio cholerae |
| ko04066 | 0.071 | 0.009 | <0.0001 | HIF-1 signaling pathway |

| | | | | |
|---|---|---|---|---|
| ko02020 | -0.183 | 0.024 | <0.0001 | Two-component system |
| ko00521 | -0.229 | 0.030 | <0.0001 | Streptomycin biosynthesis |
| ko00061 | -0.107 | 0.014 | <0.0001 | Fatty acid biosynthesis |
| ko00310 | 0.138 | 0.018 | <0.0001 | Lysine degradation |
| ko04621 | 0.050 | 0.006 | <0.0001 | NOD-like receptor signaling pathway |
| ko04212 | 0.082 | 0.011 | <0.0001 | Longevity regulating pathway - worm |
| ko05152 | 0.093 | 0.012 | <0.0001 | Tuberculosis |
| ko04214 | 0.139 | 0.018 | <0.0001 | Apoptosis - fly |
| ko01120 | -0.071 | 0.009 | <0.0001 | Microbial metabolism in diverse environments |
| ko00400 | -0.996 | 0.133 | <0.0001 | Phenylalanine, tyrosine and tryptophan biosynthesis |
| ko00010 | -0.230 | 0.031 | <0.0001 | Glycolysis / Gluconeogenesis |
| ko00740 | 0.071 | 0.010 | <0.0001 | Riboflavin metabolism |
| ko02026 | -0.450 | 0.061 | <0.0001 | Biofilm formation - Escherichia coli |
| ko00260 | -0.064 | 0.009 | <0.0001 | Glycine, serine and threonine metabolism |
| ko00920 | -0.420 | 0.057 | <0.0001 | Sulfur metabolism |
| ko04122 | -0.091 | 0.012 | <0.0001 | Sulfur relay system |
| ko00670 | 0.056 | 0.008 | <0.0001 | One carbon pool by folate |
| ko00770 | -0.202 | 0.028 | <0.0001 | Pantothenate and CoA biosynthesis |
| ko00130 | 0.106 | 0.015 | <0.0001 | Ubiquinone and other terpenoid-quinone biosynthesis |
| ko04922 | -0.345 | 0.047 | <0.0001 | Glucagon signaling pathway |
| ko00280 | 0.078 | 0.011 | <0.0001 | Valine, leucine and isoleucine degradation |
| ko01524 | 0.169 | 0.023 | <0.0001 | Platinum drug resistance |
| ko04146 | 0.113 | 0.016 | <0.0001 | Peroxisome |
| ko04070 | 0.144 | 0.020 | <0.0001 | Phosphatidylinositol signaling system |
| ko00564 | 0.060 | 0.008 | <0.0001 | Glycerophospholipid metabolism |
| ko05418 | 0.083 | 0.012 | <0.0001 | Fluid shear stress and atherosclerosis |
| ko03030 | 0.064 | 0.009 | <0.0001 | DNA replication |
| ko00630 | 0.084 | 0.012 | <0.0001 | Glyoxylate and dicarboxylate metabolism |

| | | | | |
|---|---|---|---|---|
| ko00620 | -0.315 | 0.044 | <0.0001 | Pyruvate metabolism |
| ko04016 | -0.174 | 0.025 | <0.0001 | MAPK signaling pathway - plant |
| ko01212 | -0.103 | 0.015 | <0.0001 | Fatty acid metabolism |
| ko00350 | -0.622 | 0.088 | <0.0001 | Tyrosine metabolism |
| ko03440 | 0.085 | 0.012 | <0.0001 | Homologous recombination |
| ko00195 | 0.110 | 0.016 | <0.0001 | Photosynthesis |
| ko00960 | 0.154 | 0.022 | <0.0001 | Tropane, piperidine and pyridine alkaloid biosynthesis |
| ko05204 | 0.250 | 0.036 | <0.0001 | Chemical carcinogenesis |
| ko00471 | 0.021 | 0.003 | <0.0001 | D-Glutamine and D-glutamate metabolism |
| ko00950 | 0.250 | 0.036 | <0.0001 | Isoquinoline alkaloid biosynthesis |
| ko01230 | -0.125 | 0.018 | <0.0001 | Biosynthesis of amino acids |
| ko03010 | 0.059 | 0.009 | <0.0001 | Ribosome |
| ko04013 | 0.250 | 0.036 | <0.0001 | MAPK signaling pathway - fly |
| ko04072 | 0.250 | 0.036 | <0.0001 | Phospholipase D signaling pathway |
| ko04115 | 0.250 | 0.036 | <0.0001 | p53 signaling pathway |
| ko04151 | 0.250 | 0.036 | <0.0001 | PI3K-Akt signaling pathway |
| ko04210 | 0.250 | 0.036 | <0.0001 | Apoptosis |
| ko04215 | 0.250 | 0.036 | <0.0001 | Apoptosis - multiple species |
| ko04612 | 0.250 | 0.036 | <0.0001 | Antigen processing and presentation |
| ko04657 | 0.250 | 0.036 | <0.0001 | IL-17 signaling pathway |
| ko04659 | 0.250 | 0.036 | <0.0001 | Th17 cell differentiation |
| ko04914 | 0.250 | 0.036 | <0.0001 | Progesterone-mediated oocyte maturation |
| ko04915 | 0.250 | 0.036 | <0.0001 | Estrogen signaling pathway |
| ko05120 | 0.250 | 0.036 | <0.0001 | Epithelial cell signaling in Helicobacter pylori infection |
| ko05133 | 0.250 | 0.036 | <0.0001 | Pertussis |
| ko05145 | 0.250 | 0.036 | <0.0001 | Toxoplasmosis |
| ko05161 | 0.250 | 0.036 | <0.0001 | Hepatitis B |
| ko05164 | 0.250 | 0.036 | <0.0001 | Influenza A |

| | | | | |
|---|---|---|---|---|
| ko05168 | 0.250 | 0.036 | <0.0001 | Herpes simplex infection |
| ko05210 | 0.250 | 0.036 | <0.0001 | Colorectal cancer |
| ko05215 | 0.250 | 0.036 | <0.0001 | Prostate cancer |
| ko05222 | 0.250 | 0.036 | <0.0001 | Small cell lung cancer |
| ko05231 | 0.250 | 0.036 | <0.0001 | Choline metabolism in cancer |
| ko05416 | 0.250 | 0.036 | <0.0001 | Viral myocarditis |
| ko00750 | 0.078 | 0.011 | <0.0001 | Vitamin B6 metabolism |
| ko01055 | 0.154 | 0.023 | <0.0001 | Biosynthesis of vancomycin group antibiotics |
| ko00220 | 0.059 | 0.009 | <0.0001 | Arginine biosynthesis |
| ko00520 | -0.503 | 0.074 | <0.0001 | Amino sugar and nucleotide sugar metabolism |
| ko04152 | 0.154 | 0.023 | <0.0001 | AMPK signaling pathway |
| ko02010 | -0.345 | 0.051 | <0.0001 | ABC transporters |
| ko01523 | 0.035 | 0.005 | <0.0001 | Antifolate resistance |
| ko05200 | 0.185 | 0.028 | <0.0001 | Pathways in cancer |
| ko01200 | -0.034 | 0.005 | <0.0001 | Carbon metabolism |
| ko01100 | -0.025 | 0.004 | <0.0001 | Metabolic pathways |
| ko00473 | -0.355 | 0.054 | <0.0001 | D-Alanine metabolism |
| ko04141 | 0.054 | 0.008 | <0.0001 | Protein processing in endoplasmic reticulum |
| ko05134 | 0.158 | 0.025 | <0.0001 | Legionellosis |
| ko00680 | -0.212 | 0.033 | <0.0001 | Methane metabolism |
| ko04112 | 0.129 | 0.020 | <0.0001 | Cell cycle - Caulobacter |
| ko05230 | -0.210 | 0.033 | <0.0001 | Central carbon metabolism in cancer |
| ko00785 | 0.064 | 0.010 | <0.0001 | Lipoic acid metabolism |
| ko05211 | -0.047 | 0.008 | <0.0001 | Renal cell carcinoma |
| ko00240 | 0.036 | 0.006 | <0.0001 | Pyrimidine metabolism |
| ko03060 | 0.118 | 0.019 | <0.0001 | Protein export |
| ko03018 | 0.026 | 0.004 | <0.0001 | RNA degradation |
| ko00561 | -0.248 | 0.042 | <0.0001 | Glycerolipid metabolism |

| | | | | |
|---|---|---|---|---|
| ko00270 | -0.140 | 0.024 | <0.0001 | Cysteine and methionine metabolism |
| ko02024 | -0.229 | 0.039 | <0.0001 | Quorum sensing |
| ko05010 | 0.217 | 0.037 | <0.0001 | Alzheimer's disease |
| ko00640 | -0.181 | 0.031 | <0.0001 | Propanoate metabolism |
| ko04260 | 0.250 | 0.043 | <0.0001 | Cardiac muscle contraction |
| ko01501 | -0.323 | 0.056 | <0.0001 | beta-Lactam resistance |
| ko04932 | 0.250 | 0.044 | <0.0001 | Non-alcoholic fatty liver disease (NAFLD) |
| ko05012 | 0.250 | 0.044 | <0.0001 | Parkinson's disease |
| ko00330 | -0.174 | 0.031 | <0.0001 | Arginine and proline metabolism |
| ko00710 | 0.093 | 0.017 | <0.0001 | Carbon fixation in photosynthetic organisms |
| ko03070 | 0.198 | 0.035 | <0.0001 | Bacterial secretion system |
| ko05016 | 0.250 | 0.045 | <0.0001 | Huntington's disease |
| ko01503 | -0.290 | 0.052 | <0.0001 | Cationic antimicrobial peptide (CAMP) resistance |
| ko00860 | 0.236 | 0.043 | <0.0001 | Porphyrin and chlorophyll metabolism |
| ko00190 | 0.187 | 0.034 | <0.0001 | Oxidative phosphorylation |
| ko00760 | -0.416 | 0.077 | <0.0001 | Nicotinate and nicotinamide metabolism |
| ko04940 | -0.099 | 0.019 | <0.0001 | Type I diabetes mellitus |
| ko04727 | -0.099 | 0.019 | <0.0001 | GABAergic synapse |
| ko00430 | -0.474 | 0.092 | <0.0001 | Taurine and hypotaurine metabolism |
| ko00650 | -0.174 | 0.034 | <0.0001 | Butanoate metabolism |
| ko00020 | 0.171 | 0.033 | <0.0001 | Citrate cycle (TCA cycle) |
| ko00250 | -0.019 | 0.004 | <0.0001 | Alanine, aspartate and glutamate metabolism |
| ko00460 | -0.221 | 0.044 | <0.0001 | Cyanoamino acid metabolism |
| ko00290 | -4.644 | 0.929 | <0.0001 | Valine, leucine and isoleucine biosynthesis |
| ko00030 | -0.267 | 0.054 | <0.0001 | Pentose phosphate pathway |
| ko00332 | -5.117 | 1.039 | <0.0001 | Carbapenem biosynthesis |
| ko00623 | -4.453 | 0.905 | <0.0001 | Toluene degradation |
| ko00340 | -5.107 | 1.040 | <0.0001 | Histidine metabolism |

| | | | | |
|---|---|---|---|---|
| ko00900 | 0.018 | 0.004 | <0.0001 | Terpenoid backbone biosynthesis |
| ko00603 | -4.716 | 0.969 | <0.0001 | Glycosphingolipid biosynthesis - globo and isoglobo series |
| ko00405 | -5.130 | 1.056 | <0.0001 | Phenazine biosynthesis |
| ko02025 | -5.095 | 1.050 | <0.0001 | Biofilm formation - Pseudomonas aeruginosa |
| ko03410 | -0.020 | 0.004 | <0.0001 | Base excision repair |
| ko00311 | -4.731 | 0.988 | <0.0001 | Penicillin and cephalosporin biosynthesis |
| ko00622 | -4.488 | 0.937 | <0.0001 | Xylene degradation |
| ko03430 | 0.028 | 0.006 | <0.0001 | Mismatch repair |
| ko00052 | -4.797 | 1.008 | <0.0001 | Galactose metabolism |
| ko00500 | -4.942 | 1.040 | <0.0001 | Starch and sucrose metabolism |
| ko01054 | -4.641 | 0.977 | <0.0001 | Nonribosomal peptide structures |
| ko00440 | -4.843 | 1.022 | <0.0001 | Phosphonate and phosphinate metabolism |
| ko01220 | -4.633 | 0.978 | <0.0001 | Degradation of aromatic compounds |
| ko00524 | -4.505 | 0.952 | <0.0001 | Neomycin, kanamycin and gentamicin biosynthesis |
| ko04918 | -4.684 | 1.008 | <0.0001 | Thyroid hormone synthesis |
| ko00830 | -4.656 | 1.003 | <0.0001 | Retinol metabolism |
| ko02060 | -5.105 | 1.099 | <0.0001 | Phosphotransferase system (PTS) |
| ko00621 | -4.517 | 0.977 | <0.0001 | Dioxin degradation |
| ko00362 | -4.396 | 0.952 | <0.0001 | Benzoate degradation |
| ko00072 | -4.506 | 0.979 | <0.0001 | Synthesis and degradation of ketone bodies |
| ko03320 | -4.514 | 0.982 | <0.0001 | PPAR signaling pathway |
| ko00590 | -4.520 | 0.984 | <0.0001 | Arachidonic acid metabolism |
| ko00626 | -4.618 | 1.006 | <0.0001 | Naphthalene degradation |
| ko04930 | -4.523 | 0.985 | <0.0001 | Type II diabetes mellitus |
| ko05203 | -4.523 | 0.985 | <0.0001 | Viral carcinogenesis |
| ko03013 | -4.525 | 0.988 | <0.0001 | RNA transport |
| ko00360 | -0.099 | 0.022 | <0.0001 | Phenylalanine metabolism |

| | | | | |
|---|---|---|---|---|
| ko00600 | -4.523 | 0.990 | <0.0001 | Sphingolipid metabolism |
| ko04917 | -4.529 | 0.992 | <0.0001 | Prolactin signaling pathway |
| ko00120 | -4.623 | 1.015 | <0.0001 | Primary bile acid biosynthesis |
| ko00121 | -4.623 | 1.015 | <0.0001 | Secondary bile acid biosynthesis |
| ko00625 | -4.598 | 1.011 | <0.0001 | Chloroalkane and chloroalkene degradation |
| ko00730 | -0.084 | 0.018 | <0.0001 | Thiamine metabolism |
| ko00627 | -4.390 | 0.979 | <0.0001 | Aminobenzoate degradation |
| ko00071 | -4.444 | 0.992 | <0.0001 | Fatty acid degradation |
| ko00040 | -0.332 | 0.075 | <0.0001 | Pentose and glucuronate interconversions |
| ko05150 | -4.596 | 1.032 | <0.0001 | Staphylococcus aureus infection |
| ko00511 | -4.442 | 1.002 | <0.0001 | Other glycan degradation |
| ko04080 | -4.547 | 1.037 | <0.0001 | Neuroactive ligand-receptor interaction |
| ko05166 | -4.547 | 1.037 | <0.0001 | HTLV-I infection |
| ko00361 | -4.530 | 1.037 | <0.0001 | Chlorocyclohexane and chlorobenzene degradation |
| ko00790 | -0.108 | 0.025 | <0.0001 | Folate biosynthesis |
| ko03420 | 0.021 | 0.005 | <0.0001 | Nucleotide excision repair |
| ko04138 | -4.551 | 1.060 | <0.0001 | Autophagy - yeast |
| ko05340 | -4.330 | 1.009 | <0.0001 | Primary immunodeficiency |
| ko00983 | -0.161 | 0.038 | <0.0001 | Drug metabolism - other enzymes |
| ko01210 | 0.147 | 0.035 | <0.0001 | 2-Oxocarboxylic acid metabolism |
| ko00401 | 0.064 | 0.016 | <0.0001 | Novobiocin biosynthesis |
| ko00643 | -3.866 | 0.980 | <0.0001 | Styrene degradation |
| ko00531 | -3.878 | 0.985 | <0.0001 | Glycosaminoglycan degradation |
| ko04011 | -4.559 | 1.166 | <0.0001 | MAPK signaling pathway - yeast |
| ko04142 | -3.899 | 1.006 | <0.0001 | Lysosome |
| ko04910 | -5.172 | 1.359 | <0.0001 | Insulin signaling pathway |
| ko00944 | -3.917 | 1.036 | <0.0001 | Flavone and flavonol biosynthesis |
| ko00523 | -0.139 | 0.037 | <0.0001 | Polyketide sugar unit biosynthesis |

| | | | | |
|---|---|---|---|---|
| ko00140 | -3.911 | 1.041 | <0.0001 | Steroid hormone biosynthesis |
| ko00906 | -5.185 | 1.392 | <0.0001 | Carotenoid biosynthesis |
| ko01062 | -5.189 | 1.463 | 0.001 | Biosynthesis of terpenoids and steroids |
| ko04622 | -5.211 | 1.489 | 0.001 | RIG-I-like receptor signaling pathway |
| ko01130 | -0.027 | 0.008 | 0.002 | Biosynthesis of antibiotics |
| ko00940 | 0.035 | 0.012 | 0.003 | Phenylpropanoid biosynthesis |
| ko00970 | 0.016 | 0.006 | 0.009 | Aminoacyl-tRNA biosynthesis |
| ko00450 | -0.031 | 0.012 | 0.011 | Selenocompound metabolism |
| ko00261 | 0.086 | 0.033 | 0.012 | Monobactam biosynthesis |
| ko00562 | -0.014 | 0.006 | 0.028 | Inositol phosphate metabolism |
| ko00230 | -0.006 | 0.003 | 0.046 | Purine metabolism |

**Table S14 Effect of age on functional pathway abundance for single male flies.** Predicted gene function was performed using Piphillin and the KEGG reference database (May 2017 release) using a sequence identity cut-off of 97%. DESeq2 analysis was performed to identify pathways that were differentially abundant between 11 day old and 49-day old single males. P-values are corrected for multiple testing using the Benjamini-Hochberg method.

| KEGG number | Log2 Fold Change | LFC Standard Error | Adj. *P* Value | Pathway Function |
|---|---|---|---|---|
| ko00052 | -5.400 | 1.008 | <0.0001 | Galactose metabolism |
| ko00072 | -5.139 | 0.982 | <0.0001 | Synthesis and degradation of ketone bodies |
| ko00120 | -5.374 | 1.017 | <0.0001 | Primary bile acid biosynthesis |
| ko00121 | -5.374 | 1.017 | <0.0001 | Secondary bile acid biosynthesis |
| ko00290 | -4.878 | 0.932 | <0.0001 | Valine, leucine and isoleucine biosynthesis |
| ko00311 | -5.403 | 0.992 | <0.0001 | Penicillin and cephalosporin biosynthesis |
| ko00440 | -5.503 | 1.026 | <0.0001 | Phosphonate and phosphinate metabolism |
| ko00500 | -5.487 | 1.040 | <0.0001 | Starch and sucrose metabolism |
| ko00511 | -5.254 | 1.004 | <0.0001 | Other glycan degradation |
| ko00524 | -5.143 | 0.958 | <0.0001 | Neomycin, kanamycin and gentamicin biosynthesis |
| ko00590 | -5.252 | 0.990 | <0.0001 | Arachidonic acid metabolism |
| ko00600 | -5.273 | 0.991 | <0.0001 | Sphingolipid metabolism |
| ko00603 | -5.338 | 0.974 | <0.0001 | Glycosphingolipid biosynthesis - globo and isoglobo series |
| ko00621 | -5.227 | 0.983 | <0.0001 | Dioxin degradation |
| ko00622 | -5.055 | 0.939 | <0.0001 | Xylene degradation |
| ko00623 | -4.896 | 0.910 | <0.0001 | Toluene degradation |
| ko00625 | -5.334 | 1.012 | <0.0001 | Chloroalkane and chloroalkene degradation |

| | | | | |
|---|---|---|---|---|
| ko00626 | -5.349 | 1.007 | <0.0001 | Naphthalene degradation |
| ko00627 | -5.125 | 0.981 | <0.0001 | Aminobenzoate degradation |
| ko00830 | -5.374 | 1.005 | <0.0001 | Retinol metabolism |
| ko01054 | -5.361 | 0.982 | <0.0001 | Nonribosomal peptide structures |
| ko01220 | -5.228 | 0.979 | <0.0001 | Degradation of aromatic compounds |
| ko03013 | -5.310 | 0.991 | <0.0001 | RNA transport |
| ko03320 | -5.166 | 0.984 | <0.0001 | PPAR signaling pathway |
| ko04080 | -5.455 | 1.043 | <0.0001 | Neuroactive ligand-receptor interaction |
| ko04917 | -5.312 | 0.998 | <0.0001 | Prolactin signaling pathway |
| ko04918 | -5.380 | 1.010 | <0.0001 | Thyroid hormone synthesis |
| ko04930 | -5.275 | 0.991 | <0.0001 | Type II diabetes mellitus |
| ko05166 | -5.455 | 1.043 | <0.0001 | HTLV-I infection |
| ko05203 | -5.275 | 0.991 | <0.0001 | Viral carcinogenesis |
| ko05150 | -5.367 | 1.033 | <0.0001 | Staphylococcus aureus infection |
| ko00362 | -4.933 | 0.953 | <0.0001 | Benzoate degradation |
| ko05340 | -5.213 | 1.013 | <0.0001 | Primary immunodeficiency |
| ko00071 | -5.105 | 0.993 | <0.0001 | Fatty acid degradation |
| ko00405 | -5.421 | 1.060 | <0.0001 | Phenazine biosynthesis |
| ko02060 | -5.626 | 1.099 | <0.0001 | Phosphotransferase system (PTS) |
| ko00332 | -5.321 | 1.044 | <0.0001 | Carbapenem biosynthesis |
| ko04138 | -5.430 | 1.066 | <0.0001 | Autophagy - yeast |
| ko00361 | -5.287 | 1.043 | <0.0001 | Chlorocyclohexane and chlorobenzene degradation |
| ko00340 | -5.253 | 1.041 | <0.0001 | Histidine metabolism |
| ko02025 | -5.293 | 1.053 | <0.0001 | Biofilm formation - Pseudomonas aeruginosa |
| ko00643 | -4.791 | 0.989 | <0.0001 | Styrene degradation |
| ko04142 | -4.868 | 1.011 | <0.0001 | Lysosome |
| ko00140 | -5.005 | 1.052 | <0.0001 | Steroid hormone biosynthesis |

| | | | | |
|---|---|---|---|---|
| ko00531 | -4.684 | 0.994 | <0.0001 | Glycosaminoglycan degradation |
| ko00944 | -4.918 | 1.045 | <0.0001 | Flavone and flavonol biosynthesis |
| ko04011 | -5.479 | 1.172 | <0.0001 | MAPK signaling pathway - yeast |
| ko00906 | -5.736 | 1.397 | <0.0001 | Carotenoid biosynthesis |
| ko04910 | -5.499 | 1.369 | <0.0001 | Insulin signaling pathway |
| ko01062 | -5.777 | 1.474 | <0.0001 | Biosynthesis of terpenoids and steroids |
| ko04622 | -5.874 | 1.512 | 0.001 | RIG-I-like receptor signaling pathway |
| ko00670 | 0.026 | 0.008 | 0.003 | One carbon pool by folate |
| ko04621 | 0.023 | 0.007 | 0.003 | NOD-like receptor signaling pathway |
| ko04066 | 0.031 | 0.009 | 0.004 | HIF-1 signaling pathway |
| ko01523 | 0.018 | 0.005 | 0.004 | Antifolate resistance |
| ko00564 | 0.027 | 0.008 | 0.005 | Glycerophospholipid metabolism |
| ko00400 | -0.432 | 0.133 | 0.005 | Phenylalanine, tyrosine and tryptophan biosynthesis |
| ko00740 | 0.031 | 0.010 | 0.005 | Riboflavin metabolism |
| ko04212 | 0.035 | 0.011 | 0.005 | Longevity regulating pathway - worm |
| ko00051 | -0.245 | 0.077 | 0.005 | Fructose and mannose metabolism |
| ko00220 | 0.028 | 0.009 | 0.005 | Arginine biosynthesis |
| ko00660 | 0.038 | 0.012 | 0.005 | C5-Branched dibasic acid metabolism |
| ko00908 | 0.037 | 0.012 | 0.005 | Zeatin biosynthesis |
| ko03008 | 0.037 | 0.012 | 0.005 | Ribosome biogenesis in eukaryotes |
| ko03030 | 0.028 | 0.009 | 0.005 | DNA replication |
| ko04213 | 0.038 | 0.012 | 0.005 | Longevity regulating pathway - multiple species |
| ko04626 | 0.038 | 0.012 | 0.005 | Plant-pathogen interaction |
| ko04724 | 0.037 | 0.012 | 0.005 | Glutamatergic synapse |
| ko05111 | -0.213 | 0.067 | 0.005 | Biofilm formation - Vibrio cholerae |
| ko05205 | 0.037 | 0.012 | 0.005 | Proteoglycans in cancer |
| ko05418 | 0.037 | 0.012 | 0.005 | Fluid shear stress and atherosclerosis |

| | | | | |
|---|---|---|---|---|
| ko04068 | 0.037 | 0.012 | 0.005 | FoxO signaling pathway |
| ko04211 | 0.037 | 0.012 | 0.005 | Longevity regulating pathway |
| ko05152 | 0.039 | 0.012 | 0.005 | Tuberculosis |
| ko05014 | 0.037 | 0.012 | 0.005 | Amyotrophic lateral sclerosis (ALS) |
| ko00750 | 0.036 | 0.012 | 0.006 | Vitamin B6 metabolism |
| ko03010 | 0.026 | 0.009 | 0.007 | Ribosome |
| ko00630 | 0.036 | 0.012 | 0.007 | Glyoxylate and dicarboxylate metabolism |
| ko03018 | 0.014 | 0.004 | 0.007 | RNA degradation |
| ko04141 | 0.027 | 0.009 | 0.007 | Protein processing in endoplasmic reticulum |
| ko00350 | -0.267 | 0.088 | 0.008 | Tyrosine metabolism |
| ko04146 | 0.047 | 0.016 | 0.008 | Peroxisome |
| ko00130 | 0.043 | 0.015 | 0.009 | Ubiquinone and other terpenoid-quinone biosynthesis |
| ko03440 | 0.036 | 0.012 | 0.009 | Homologous recombination |
| ko00240 | 0.017 | 0.006 | 0.009 | Pyrimidine metabolism |
| ko00280 | 0.032 | 0.011 | 0.009 | Valine, leucine and isoleucine degradation |
| ko00310 | 0.053 | 0.018 | 0.009 | Lysine degradation |
| ko00980 | -0.043 | 0.014 | 0.009 | Metabolism of xenobiotics by cytochrome P450 |
| ko00982 | -0.043 | 0.014 | 0.009 | Drug metabolism - cytochrome P450 |
| ko01040 | -0.042 | 0.014 | 0.009 | Biosynthesis of unsaturated fatty acids |
| ko04214 | 0.054 | 0.018 | 0.009 | Apoptosis - fly |
| ko04070 | 0.058 | 0.020 | 0.010 | Phosphatidylinositol signaling system |
| ko00960 | 0.064 | 0.022 | 0.011 | Tropane, piperidine and pyridine alkaloid biosynthesis |
| ko00195 | 0.045 | 0.016 | 0.011 | Photosynthesis |
| ko00910 | -0.058 | 0.020 | 0.011 | Nitrogen metabolism |
| ko04922 | -0.135 | 0.047 | 0.011 | Glucagon signaling pathway |
| ko01524 | 0.067 | 0.023 | 0.011 | Platinum drug resistance |
| ko00520 | -0.210 | 0.074 | 0.012 | Amino sugar and nucleotide sugar metabolism |

| | | | | |
|---|---|---|---|---|
| ko01055 | 0.064 | 0.023 | 0.012 | Biosynthesis of vancomycin group antibiotics |
| ko04152 | 0.064 | 0.023 | 0.013 | AMPK signaling pathway |
| ko02026 | -0.170 | 0.061 | 0.013 | Biofilm formation - Escherichia coli |
| ko00620 | -0.122 | 0.044 | 0.013 | Pyruvate metabolism |
| ko00900 | 0.010 | 0.004 | 0.013 | Terpenoid backbone biosynthesis |
| ko00920 | -0.158 | 0.057 | 0.014 | Sulfur metabolism |
| ko00010 | -0.085 | 0.031 | 0.015 | Glycolysis / Gluconeogenesis |
| ko02020 | -0.064 | 0.024 | 0.015 | Two-component system |
| ko00521 | -0.081 | 0.030 | 0.015 | Streptomycin biosynthesis |
| ko00770 | -0.075 | 0.028 | 0.015 | Pantothenate and CoA biosynthesis |
| ko04016 | -0.067 | 0.025 | 0.015 | MAPK signaling pathway - plant |
| ko00473 | -0.145 | 0.054 | 0.017 | D-Alanine metabolism |
| ko00950 | 0.096 | 0.037 | 0.017 | Isoquinoline alkaloid biosynthesis |
| ko02010 | -0.137 | 0.051 | 0.017 | ABC transporters |
| ko04013 | 0.096 | 0.037 | 0.017 | MAPK signaling pathway - fly |
| ko04072 | 0.096 | 0.037 | 0.017 | Phospholipase D signaling pathway |
| ko04115 | 0.096 | 0.037 | 0.017 | p53 signaling pathway |
| ko04151 | 0.096 | 0.037 | 0.017 | PI3K-Akt signaling pathway |
| ko04210 | 0.096 | 0.037 | 0.017 | Apoptosis |
| ko04215 | 0.096 | 0.037 | 0.017 | Apoptosis - multiple species |
| ko04612 | 0.096 | 0.037 | 0.017 | Antigen processing and presentation |
| ko04657 | 0.096 | 0.037 | 0.017 | IL-17 signaling pathway |
| ko04659 | 0.096 | 0.037 | 0.017 | Th17 cell differentiation |
| ko04914 | 0.096 | 0.037 | 0.017 | Progesterone-mediated oocyte maturation |
| ko04915 | 0.096 | 0.037 | 0.017 | Estrogen signaling pathway |
| ko05120 | 0.096 | 0.037 | 0.017 | Epithelial cell signaling in Helicobacter pylori infection |
| ko05133 | 0.096 | 0.037 | 0.017 | Pertussis |

| | | | | |
|---|---|---|---|---|
| ko05145 | 0.096 | 0.037 | 0.017 | Toxoplasmosis |
| ko05161 | 0.096 | 0.037 | 0.017 | Hepatitis B |
| ko05164 | 0.096 | 0.037 | 0.017 | Influenza A |
| ko05168 | 0.096 | 0.037 | 0.017 | Herpes simplex infection |
| ko05200 | 0.073 | 0.028 | 0.017 | Pathways in cancer |
| ko05204 | 0.095 | 0.036 | 0.017 | Chemical carcinogenesis |
| ko05210 | 0.096 | 0.037 | 0.017 | Colorectal cancer |
| ko05215 | 0.096 | 0.037 | 0.017 | Prostate cancer |
| ko05222 | 0.096 | 0.037 | 0.017 | Small cell lung cancer |
| ko05231 | 0.096 | 0.037 | 0.017 | Choline metabolism in cancer |
| ko05416 | 0.096 | 0.037 | 0.017 | Viral myocarditis |
| ko00061 | -0.036 | 0.014 | 0.017 | Fatty acid biosynthesis |
| ko04112 | 0.053 | 0.020 | 0.018 | Cell cycle - Caulobacter |
| ko05134 | 0.063 | 0.025 | 0.018 | Legionellosis |
| ko03060 | 0.049 | 0.019 | 0.021 | Protein export |
| ko00710 | 0.041 | 0.017 | 0.025 | Carbon fixation in photosynthetic organisms |
| ko01212 | -0.036 | 0.015 | 0.026 | Fatty acid metabolism |
| ko00680 | -0.081 | 0.033 | 0.026 | Methane metabolism |
| ko05230 | -0.081 | 0.033 | 0.027 | Central carbon metabolism in cancer |
| ko00785 | 0.025 | 0.011 | 0.027 | Lipoic acid metabolism |
| ko00471 | 0.008 | 0.003 | 0.031 | D-Glutamine and D-glutamate metabolism |
| ko01501 | -0.132 | 0.056 | 0.032 | beta-Lactam resistance |
| ko03430 | 0.014 | 0.006 | 0.034 | Mismatch repair |
| ko00561 | -0.098 | 0.042 | 0.034 | Glycerolipid metabolism |
| ko00760 | -0.178 | 0.077 | 0.036 | Nicotinate and nicotinamide metabolism |
| ko00430 | -0.210 | 0.092 | 0.038 | Taurine and hypotaurine metabolism |
| ko01120 | -0.021 | 0.009 | 0.038 | Microbial metabolism in diverse environments |
| ko02024 | -0.089 | 0.039 | 0.038 | Quorum sensing |

| | | | | |
|---|---|---|---|---|
| ko05010 | 0.085 | 0.037 | 0.038 | Alzheimer's disease |
| ko01503 | -0.118 | 0.052 | 0.038 | Cationic antimicrobial peptide (CAMP) resistance |
| ko00640 | -0.069 | 0.031 | 0.043 | Propanoate metabolism |
| ko04260 | 0.096 | 0.043 | 0.043 | Cardiac muscle contraction |
| ko03070 | 0.078 | 0.035 | 0.045 | Bacterial secretion system |
| ko00330 | -0.068 | 0.031 | 0.045 | Arginine and proline metabolism |
| ko03420 | 0.011 | 0.005 | 0.045 | Nucleotide excision repair |
| ko01230 | -0.040 | 0.018 | 0.046 | Biosynthesis of amino acids |
| ko04122 | -0.027 | 0.012 | 0.046 | Sulfur relay system |
| ko04932 | 0.096 | 0.044 | 0.046 | Non-alcoholic fatty liver disease (NAFLD) |
| ko05012 | 0.096 | 0.044 | 0.046 | Parkinson's disease |
| ko00401 | 0.034 | 0.016 | 0.046 | Novobiocin biosynthesis |
| ko00190 | 0.074 | 0.034 | 0.048 | Oxidative phosphorylation |
| ko05016 | 0.096 | 0.045 | 0.049 | Huntington's disease |

**Table S15 Effect of age on functional pathway abundance for grouped male flies.** Predicted gene function was performed using Piphillin and the KEGG reference database (May 2017 release) using a sequence identity cut-off of 97%. DESeq2 analysis was performed to identify pathways that were differentially abundant between 11 day old and 49-day old grouped (co-aged) males. P-values are corrected for multiple testing using the Benjamini-Hochberg method.

| KEGG number | Log2 Fold Change | LFC Standard Error | Adj. *P* Value | Pathway Function |
|---|---|---|---|---|
| ko00908 | 0.101 | 0.012 | <0.0001 | Zeatin biosynthesis |
| ko01040 | -0.124 | 0.014 | <0.0001 | Biosynthesis of unsaturated fatty acids |
| ko03008 | 0.101 | 0.012 | <0.0001 | Ribosome biogenesis in eukaryotes |
| ko04068 | 0.101 | 0.012 | <0.0001 | FoxO signaling pathway |
| ko04211 | 0.101 | 0.012 | <0.0001 | Longevity regulating pathway |
| ko04724 | 0.101 | 0.011 | <0.0001 | Glutamatergic synapse |
| ko05014 | 0.101 | 0.012 | <0.0001 | Amyotrophic lateral sclerosis (ALS) |
| ko05205 | 0.101 | 0.012 | <0.0001 | Proteoglycans in cancer |
| ko00980 | -0.124 | 0.014 | <0.0001 | Metabolism of xenobiotics by cytochrome P450 |
| ko00982 | -0.124 | 0.014 | <0.0001 | Drug metabolism - cytochrome P450 |
| ko00660 | 0.101 | 0.012 | <0.0001 | C5-Branched dibasic acid metabolism |
| ko04621 | 0.057 | 0.007 | <0.0001 | NOD-like receptor signaling pathway |
| ko04626 | 0.101 | 0.012 | <0.0001 | Plant-pathogen interaction |
| ko04066 | 0.079 | 0.009 | <0.0001 | HIF-1 signaling pathway |
| ko04213 | 0.101 | 0.012 | <0.0001 | Longevity regulating pathway - multiple species |
| ko00471 | 0.027 | 0.003 | <0.0001 | D-Glutamine and D-glutamate metabolism |
| ko04212 | 0.090 | 0.011 | <0.0001 | Longevity regulating pathway - worm |
| ko05152 | 0.101 | 0.012 | <0.0001 | Tuberculosis |
| ko00670 | 0.063 | 0.008 | <0.0001 | One carbon pool by folate |

| | | | | |
|---|---|---|---|---|
| ko00740 | 0.079 | 0.010 | <0.0001 | Riboflavin metabolism |
| ko00310 | 0.147 | 0.018 | <0.0001 | Lysine degradation |
| ko00051 | -0.619 | 0.077 | <0.0001 | Fructose and mannose metabolism |
| ko04214 | 0.147 | 0.018 | <0.0001 | Apoptosis - fly |
| ko00910 | -0.162 | 0.020 | <0.0001 | Nitrogen metabolism |
| ko00564 | 0.067 | 0.008 | <0.0001 | Glycerophospholipid metabolism |
| ko05111 | -0.537 | 0.067 | <0.0001 | Biofilm formation - Vibrio cholerae |
| ko03030 | 0.071 | 0.009 | <0.0001 | DNA replication |
| ko05418 | 0.092 | 0.012 | <0.0001 | Fluid shear stress and atherosclerosis |
| ko00280 | 0.085 | 0.011 | <0.0001 | Valine, leucine and isoleucine degradation |
| ko00130 | 0.114 | 0.015 | <0.0001 | Ubiquinone and other terpenoid-quinone biosynthesis |
| ko00630 | 0.092 | 0.012 | <0.0001 | Glyoxylate and dicarboxylate metabolism |
| ko01523 | 0.042 | 0.005 | <0.0001 | Antifolate resistance |
| ko04146 | 0.122 | 0.016 | <0.0001 | Peroxisome |
| ko02020 | -0.182 | 0.024 | <0.0001 | Two-component system |
| ko00521 | -0.229 | 0.030 | <0.0001 | Streptomycin biosynthesis |
| ko04070 | 0.153 | 0.020 | <0.0001 | Phosphatidylinositol signaling system |
| ko01524 | 0.180 | 0.023 | <0.0001 | Platinum drug resistance |
| ko03010 | 0.066 | 0.009 | <0.0001 | Ribosome |
| ko00400 | -1.018 | 0.133 | <0.0001 | Phenylalanine, tyrosine and tryptophan biosynthesis |
| ko00220 | 0.066 | 0.009 | <0.0001 | Arginine biosynthesis |
| ko03440 | 0.093 | 0.012 | <0.0001 | Homologous recombination |
| ko00061 | -0.105 | 0.014 | <0.0001 | Fatty acid biosynthesis |
| ko00750 | 0.087 | 0.012 | <0.0001 | Vitamin B6 metabolism |
| ko00010 | -0.232 | 0.031 | <0.0001 | Glycolysis / Gluconeogenesis |
| ko02026 | -0.455 | 0.061 | <0.0001 | Biofilm formation - Escherichia coli |
| ko00195 | 0.118 | 0.016 | <0.0001 | Photosynthesis |
| ko00920 | -0.424 | 0.057 | <0.0001 | Sulfur metabolism |

| | | | | |
|---|---|---|---|---|
| ko04922 | -0.351 | 0.047 | <0.0001 | Glucagon signaling pathway |
| ko00960 | 0.165 | 0.022 | <0.0001 | Tropane, piperidine and pyridine alkaloid biosynthesis |
| ko00770 | -0.204 | 0.028 | <0.0001 | Pantothenate and CoA biosynthesis |
| ko00620 | -0.320 | 0.044 | <0.0001 | Pyruvate metabolism |
| ko01055 | 0.165 | 0.023 | <0.0001 | Biosynthesis of vancomycin group antibiotics |
| ko01120 | -0.068 | 0.009 | <0.0001 | Microbial metabolism in diverse environments |
| ko05204 | 0.263 | 0.036 | <0.0001 | Chemical carcinogenesis |
| ko00350 | -0.641 | 0.088 | <0.0001 | Tyrosine metabolism |
| ko00950 | 0.263 | 0.037 | <0.0001 | Isoquinoline alkaloid biosynthesis |
| ko04013 | 0.263 | 0.037 | <0.0001 | MAPK signaling pathway - fly |
| ko04072 | 0.263 | 0.037 | <0.0001 | Phospholipase D signaling pathway |
| ko04115 | 0.263 | 0.037 | <0.0001 | p53 signaling pathway |
| ko04141 | 0.062 | 0.009 | <0.0001 | Protein processing in endoplasmic reticulum |
| ko04151 | 0.263 | 0.037 | <0.0001 | PI3K-Akt signaling pathway |
| ko04152 | 0.165 | 0.023 | <0.0001 | AMPK signaling pathway |
| ko04210 | 0.263 | 0.037 | <0.0001 | Apoptosis |
| ko04215 | 0.263 | 0.037 | <0.0001 | Apoptosis - multiple species |
| ko04612 | 0.263 | 0.037 | <0.0001 | Antigen processing and presentation |
| ko04657 | 0.263 | 0.037 | <0.0001 | IL-17 signaling pathway |
| ko04659 | 0.263 | 0.037 | <0.0001 | Th17 cell differentiation |
| ko04914 | 0.263 | 0.037 | <0.0001 | Progesterone-mediated oocyte maturation |
| ko04915 | 0.263 | 0.037 | <0.0001 | Estrogen signaling pathway |
| ko05120 | 0.263 | 0.037 | <0.0001 | Epithelial cell signaling in Helicobacter pylori infection |
| ko05133 | 0.263 | 0.037 | <0.0001 | Pertussis |
| ko05145 | 0.263 | 0.037 | <0.0001 | Toxoplasmosis |
| ko05161 | 0.263 | 0.037 | <0.0001 | Hepatitis B |
| ko05164 | 0.263 | 0.037 | <0.0001 | Influenza A |
| ko05168 | 0.263 | 0.037 | <0.0001 | Herpes simplex infection |

| | | | | |
|---|---|---|---|---|
| ko05210 | 0.263 | 0.037 | <0.0001 | Colorectal cancer |
| ko05215 | 0.263 | 0.037 | <0.0001 | Prostate cancer |
| ko05222 | 0.263 | 0.037 | <0.0001 | Small cell lung cancer |
| ko05231 | 0.263 | 0.037 | <0.0001 | Choline metabolism in cancer |
| ko05416 | 0.263 | 0.037 | <0.0001 | Viral myocarditis |
| ko03018 | 0.032 | 0.004 | <0.0001 | RNA degradation |
| ko00623 | -6.513 | 0.910 | <0.0001 | Toluene degradation |
| ko04016 | -0.176 | 0.025 | <0.0001 | MAPK signaling pathway - plant |
| ko00290 | -6.632 | 0.933 | <0.0001 | Valine, leucine and isoleucine biosynthesis |
| ko00240 | 0.042 | 0.006 | <0.0001 | Pyrimidine metabolism |
| ko05200 | 0.196 | 0.028 | <0.0001 | Pathways in cancer |
| ko04122 | -0.088 | 0.012 | <0.0001 | Sulfur relay system |
| ko00622 | -6.632 | 0.939 | <0.0001 | Xylene degradation |
| ko00603 | -6.798 | 0.973 | <0.0001 | Glycosphingolipid biosynthesis - globo and isoglobo series |
| ko00520 | -0.517 | 0.074 | <0.0001 | Amino sugar and nucleotide sugar metabolism |
| ko00524 | -6.678 | 0.957 | <0.0001 | Neomycin, kanamycin and gentamicin biosynthesis |
| ko01212 | -0.102 | 0.015 | <0.0001 | Fatty acid metabolism |
| ko01054 | -6.827 | 0.982 | <0.0001 | Nonribosomal peptide structures |
| ko00362 | -6.616 | 0.953 | <0.0001 | Benzoate degradation |
| ko00260 | -0.061 | 0.009 | <0.0001 | Glycine, serine and threonine metabolism |
| ko00311 | -6.876 | 0.991 | <0.0001 | Penicillin and cephalosporin biosynthesis |
| ko01220 | -6.776 | 0.979 | <0.0001 | Degradation of aromatic compounds |
| ko00621 | -6.765 | 0.983 | <0.0001 | Dioxin degradation |
| ko00072 | -6.744 | 0.982 | <0.0001 | Synthesis and degradation of ketone bodies |
| ko03320 | -6.761 | 0.984 | <0.0001 | PPAR signaling pathway |
| ko00480 | -0.030 | 0.004 | <0.0001 | Glutathione metabolism |
| ko02010 | -0.352 | 0.051 | <0.0001 | ABC transporters |
| ko04930 | -6.782 | 0.991 | <0.0001 | Type II diabetes mellitus |

| | | | | |
|---|---|---|---|---|
| ko05203 | -6.782 | 0.991 | <0.0001 | Viral carcinogenesis |
| ko05134 | 0.168 | 0.025 | <0.0001 | Legionellosis |
| ko03013 | -6.769 | 0.991 | <0.0001 | RNA transport |
| ko00052 | -6.875 | 1.008 | <0.0001 | Galactose metabolism |
| ko00627 | -6.690 | 0.981 | <0.0001 | Aminobenzoate degradation |
| ko00830 | -6.852 | 1.005 | <0.0001 | Retinol metabolism |
| ko00590 | -6.741 | 0.989 | <0.0001 | Arachidonic acid metabolism |
| ko00600 | -6.750 | 0.991 | <0.0001 | Sphingolipid metabolism |
| ko04917 | -6.782 | 0.997 | <0.0001 | Prolactin signaling pathway |
| ko04918 | -6.850 | 1.009 | <0.0001 | Thyroid hormone synthesis |
| ko00440 | -6.950 | 1.025 | <0.0001 | Phosphonate and phosphinate metabolism |
| ko00626 | -6.826 | 1.007 | <0.0001 | Naphthalene degradation |
| ko04112 | 0.138 | 0.020 | <0.0001 | Cell cycle - Caulobacter |
| ko00071 | -6.723 | 0.993 | <0.0001 | Fatty acid degradation |
| ko00625 | -6.823 | 1.012 | <0.0001 | Chloroalkane and chloroalkene degradation |
| ko00120 | -6.847 | 1.017 | <0.0001 | Primary bile acid biosynthesis |
| ko00121 | -6.847 | 1.017 | <0.0001 | Secondary bile acid biosynthesis |
| ko01230 | -0.122 | 0.018 | <0.0001 | Biosynthesis of amino acids |
| ko00785 | 0.070 | 0.010 | <0.0001 | Lipoic acid metabolism |
| ko00511 | -6.744 | 1.004 | <0.0001 | Other glycan degradation |
| ko00473 | -0.364 | 0.054 | <0.0001 | D-Alanine metabolism |
| ko00500 | -6.958 | 1.040 | <0.0001 | Starch and sucrose metabolism |
| ko05340 | -6.724 | 1.013 | <0.0001 | Primary immunodeficiency |
| ko05150 | -6.848 | 1.033 | <0.0001 | Staphylococcus aureus infection |
| ko04080 | -6.897 | 1.042 | <0.0001 | Neuroactive ligand-receptor interaction |
| ko05166 | -6.897 | 1.042 | <0.0001 | HTLV-I infection |
| ko00332 | -6.872 | 1.043 | <0.0001 | Carbapenem biosynthesis |
| ko00340 | -6.843 | 1.041 | <0.0001 | Histidine metabolism |

| | | | | |
|---|---|---|---|---|
| ko00361 | -6.824 | 1.042 | <0.0001 | Chlorocyclohexane and chlorobenzene degradation |
| ko03060 | 0.127 | 0.019 | <0.0001 | Protein export |
| ko00405 | -6.909 | 1.059 | <0.0001 | Phenazine biosynthesis |
| ko04138 | -6.915 | 1.065 | <0.0001 | Autophagy - yeast |
| ko00680 | -0.215 | 0.033 | <0.0001 | Methane metabolism |
| ko02025 | -6.757 | 1.052 | <0.0001 | Biofilm formation - Pseudomonas aeruginosa |
| ko00643 | -6.352 | 0.990 | <0.0001 | Styrene degradation |
| ko02060 | -7.050 | 1.099 | <0.0001 | Phosphotransferase system (PTS) |
| ko05230 | -0.213 | 0.033 | <0.0001 | Central carbon metabolism in cancer |
| ko00531 | -6.264 | 0.995 | <0.0001 | Glycosaminoglycan degradation |
| ko00900 | 0.023 | 0.004 | <0.0001 | Terpenoid backbone biosynthesis |
| ko04142 | -6.357 | 1.011 | <0.0001 | Lysosome |
| ko00944 | -6.449 | 1.046 | <0.0001 | Flavone and flavonol biosynthesis |
| ko05010 | 0.229 | 0.037 | <0.0001 | Alzheimer's disease |
| ko00140 | -6.455 | 1.051 | <0.0001 | Steroid hormone biosynthesis |
| ko00710 | 0.102 | 0.017 | <0.0001 | Carbon fixation in photosynthetic organisms |
| ko04260 | 0.263 | 0.043 | <0.0001 | Cardiac muscle contraction |
| ko00561 | -0.252 | 0.042 | <0.0001 | Glycerolipid metabolism |
| ko04932 | 0.263 | 0.044 | <0.0001 | Non-alcoholic fatty liver disease (NAFLD) |
| ko05012 | 0.263 | 0.044 | <0.0001 | Parkinson's disease |
| ko02024 | -0.232 | 0.039 | <0.0001 | Quorum sensing |
| ko04011 | -6.922 | 1.171 | <0.0001 | MAPK signaling pathway - yeast |
| ko03070 | 0.209 | 0.035 | <0.0001 | Bacterial secretion system |
| ko01501 | -0.331 | 0.056 | <0.0001 | beta-Lactam resistance |
| ko00640 | -0.183 | 0.031 | <0.0001 | Propanoate metabolism |
| ko01200 | -0.030 | 0.005 | <0.0001 | Carbon metabolism |
| ko05016 | 0.263 | 0.045 | <0.0001 | Huntington's disease |
| ko00860 | 0.249 | 0.043 | <0.0001 | Porphyrin and chlorophyll metabolism |

| | | | | |
|---|---|---|---|---|
| ko00190 | 0.198 | 0.034 | <0.0001 | Oxidative phosphorylation |
| ko00270 | -0.137 | 0.024 | <0.0001 | Cysteine and methionine metabolism |
| ko00330 | -0.177 | 0.031 | <0.0001 | Arginine and proline metabolism |
| ko01503 | -0.297 | 0.052 | <0.0001 | Cationic antimicrobial peptide (CAMP) resistance |
| ko03430 | 0.034 | 0.006 | <0.0001 | Mismatch repair |
| ko00760 | -0.429 | 0.077 | <0.0001 | Nicotinate and nicotinamide metabolism |
| ko05211 | -0.044 | 0.008 | <0.0001 | Renal cell carcinoma |
| ko00020 | 0.181 | 0.033 | <0.0001 | Citrate cycle (TCA cycle) |
| ko03420 | 0.027 | 0.005 | <0.0001 | Nucleotide excision repair |
| ko01100 | -0.020 | 0.004 | <0.0001 | Metabolic pathways |
| ko00430 | -0.492 | 0.092 | <0.0001 | Taurine and hypotaurine metabolism |
| ko04940 | -0.099 | 0.019 | <0.0001 | Type I diabetes mellitus |
| ko04727 | -0.100 | 0.019 | <0.0001 | GABAergic synapse |
| ko00650 | -0.176 | 0.034 | <0.0001 | Butanoate metabolism |
| ko04910 | -7.078 | 1.367 | <0.0001 | Insulin signaling pathway |
| ko00460 | -0.227 | 0.044 | <0.0001 | Cyanoamino acid metabolism |
| ko00906 | -7.144 | 1.396 | <0.0001 | Carotenoid biosynthesis |
| ko00030 | -0.274 | 0.054 | <0.0001 | Pentose phosphate pathway |
| ko01062 | -7.165 | 1.471 | <0.0001 | Biosynthesis of terpenoids and steroids |
| ko04622 | -7.230 | 1.505 | <0.0001 | RIG-I-like receptor signaling pathway |
| ko00401 | 0.073 | 0.016 | <0.0001 | Novobiocin biosynthesis |
| ko00040 | -0.344 | 0.075 | <0.0001 | Pentose and glucuronate interconversions |
| ko00360 | -0.100 | 0.022 | <0.0001 | Phenylalanine metabolism |
| ko01210 | 0.157 | 0.035 | <0.0001 | 2-Oxocarboxylic acid metabolism |
| ko00730 | -0.083 | 0.018 | <0.0001 | Thiamine metabolism |
| ko00983 | -0.164 | 0.038 | <0.0001 | Drug metabolism - other enzymes |
| ko00790 | -0.103 | 0.025 | <0.0001 | Folate biosynthesis |
| ko03410 | -0.016 | 0.004 | <0.0001 | Base excision repair |

| | | | | |
|---|---|---|---|---|
| ko00250 | -0.014 | 0.004 | <0.0001 | Alanine, aspartate and glutamate metabolism |
| ko00523 | -0.134 | 0.037 | <0.0001 | Polyketide sugar unit biosynthesis |
| ko00970 | 0.021 | 0.006 | <0.0001 | Aminoacyl-tRNA biosynthesis |
| ko00780 | 0.007 | 0.002 | 0.001 | Biotin metabolism |
| ko00940 | 0.040 | 0.012 | 0.001 | Phenylpropanoid biosynthesis |
| ko01051 | 0.014 | 0.005 | 0.003 | Biosynthesis of ansamycins |
| ko04931 | 0.014 | 0.005 | 0.003 | Insulin resistance |
| ko00261 | 0.095 | 0.033 | 0.005 | Monobactam biosynthesis |
| ko00903 | -2.574 | 0.960 | 0.010 | Limonene and pinene degradation |
| ko00791 | -3.575 | 1.352 | 0.011 | Atrazine degradation |
| ko01130 | -0.022 | 0.008 | 0.012 | Biosynthesis of antibiotics |
| ko01502 | 0.005 | 0.002 | 0.020 | Vancomycin resistance |
| ko00720 | 0.011 | 0.005 | 0.026 | Carbon fixation pathways in prokaryotes |
| ko00364 | -2.897 | 1.244 | 0.026 | Fluorobenzoate degradation |
| ko00450 | -0.025 | 0.012 | 0.043 | Selenocompound metabolism |
| ko00300 | 0.029 | 0.014 | 0.045 | Lysine biosynthesis |
| ko04216 | -2.708 | 1.306 | 0.049 | Ferroptosis |

**Table S16 Effect of grouping on functional pathway abundance for male flies for co-aged groups.** Predicted gene function was performed using Piphillin and the KEGG reference database (May 2017 release) using a sequence identity cut-off of 97%. DESeq2 analysis was performed to identify pathways that were differentially abundant between 49-day old single males and males kept in co-aged groups of 10. P-values are corrected for multiple testing using the Benjamini-Hochberg method.

| KEGG number | Log2 Fold Change | LFC Standard Error | Adj. *P* Value | Pathway Function |
|---|---|---|---|---|
| ko00908 | 0.064 | 0.012 | <0.0001 | Zeatin biosynthesis |
| ko00980 | -0.081 | 0.014 | <0.0001 | Metabolism of xenobiotics by cytochrome P450 |
| ko00982 | -0.081 | 0.014 | <0.0001 | Drug metabolism - cytochrome P450 |
| ko01040 | -0.082 | 0.014 | <0.0001 | Biosynthesis of unsaturated fatty acids |
| ko03008 | 0.064 | 0.012 | <0.0001 | Ribosome biogenesis in eukaryotes |
| ko04068 | 0.064 | 0.012 | <0.0001 | FoxO signaling pathway |
| ko04211 | 0.064 | 0.012 | <0.0001 | Longevity regulating pathway |
| ko04724 | 0.064 | 0.012 | <0.0001 | Glutamatergic synapse |
| ko05014 | 0.064 | 0.012 | <0.0001 | Amyotrophic lateral sclerosis (ALS) |
| ko05205 | 0.064 | 0.012 | <0.0001 | Proteoglycans in cancer |
| ko00471 | 0.018 | 0.003 | <0.0001 | D-Glutamine and D-glutamate metabolism |
| ko00660 | 0.063 | 0.012 | <0.0001 | C5-Branched dibasic acid metabolism |
| ko04626 | 0.063 | 0.012 | <0.0001 | Plant-pathogen interaction |
| ko04213 | 0.063 | 0.012 | <0.0001 | Longevity regulating pathway - multiple species |
| ko00310 | 0.094 | 0.018 | <0.0001 | Lysine degradation |
| ko00910 | -0.105 | 0.020 | <0.0001 | Nitrogen metabolism |
| ko04066 | 0.048 | 0.009 | <0.0001 | HIF-1 signaling pathway |
| ko04214 | 0.093 | 0.018 | <0.0001 | Apoptosis - fly |
| ko04621 | 0.034 | 0.007 | <0.0001 | NOD-like receptor signaling pathway |
| ko04212 | 0.055 | 0.011 | <0.0001 | Longevity regulating pathway - worm |
| ko05152 | 0.062 | 0.012 | <0.0001 | Tuberculosis |

| | | | | |
|---|---|---|---|---|
| ko00480 | -0.022 | 0.004 | <0.0001 | Glutathione metabolism |
| ko02020 | -0.118 | 0.024 | <0.0001 | Two-component system |
| ko00521 | -0.148 | 0.030 | <0.0001 | Streptomycin biosynthesis |
| ko01120 | -0.046 | 0.009 | <0.0001 | Microbial metabolism in diverse environments |
| ko00061 | -0.069 | 0.014 | <0.0001 | Fatty acid biosynthesis |
| ko00280 | 0.053 | 0.011 | <0.0001 | Valine, leucine and isoleucine degradation |
| ko00051 | -0.374 | 0.077 | <0.0001 | Fructose and mannose metabolism |
| ko00740 | 0.047 | 0.010 | <0.0001 | Riboflavin metabolism |
| ko04122 | -0.061 | 0.012 | <0.0001 | Sulfur relay system |
| ko00130 | 0.071 | 0.015 | <0.0001 | Ubiquinone and other terpenoid-quinone biosynthesis |
| ko00260 | -0.043 | 0.009 | <0.0001 | Glycine, serine and threonine metabolism |
| ko05111 | -0.323 | 0.067 | <0.0001 | Biofilm formation - Vibrio cholerae |
| ko00670 | 0.037 | 0.008 | <0.0001 | One carbon pool by folate |
| ko01524 | 0.113 | 0.023 | <0.0001 | Platinum drug resistance |
| ko03030 | 0.043 | 0.009 | <0.0001 | DNA replication |
| ko04070 | 0.095 | 0.020 | <0.0001 | Phosphatidylinositol signaling system |
| ko00564 | 0.040 | 0.008 | <0.0001 | Glycerophospholipid metabolism |
| ko00630 | 0.056 | 0.012 | <0.0001 | Glyoxylate and dicarboxylate metabolism |
| ko04146 | 0.075 | 0.016 | <0.0001 | Peroxisome |
| ko00010 | -0.146 | 0.031 | <0.0001 | Glycolysis / Gluconeogenesis |
| ko05418 | 0.055 | 0.012 | <0.0001 | Fluid shear stress and atherosclerosis |
| ko02026 | -0.285 | 0.061 | <0.0001 | Biofilm formation - Escherichia coli |
| ko00195 | 0.073 | 0.016 | <0.0001 | Photosynthesis |
| ko00770 | -0.129 | 0.028 | <0.0001 | Pantothenate and CoA biosynthesis |
| ko00920 | -0.266 | 0.057 | <0.0001 | Sulfur metabolism |
| ko00950 | 0.167 | 0.037 | <0.0001 | Isoquinoline alkaloid biosynthesis |
| ko03010 | 0.039 | 0.009 | <0.0001 | Ribosome |
| ko03440 | 0.056 | 0.012 | <0.0001 | Homologous recombination |
| ko04013 | 0.167 | 0.037 | <0.0001 | MAPK signaling pathway - fly |
| ko04072 | 0.167 | 0.037 | <0.0001 | Phospholipase D signaling pathway |
| ko04115 | 0.167 | 0.037 | <0.0001 | p53 signaling pathway |

| | | | | |
|---|---|---|---|---|
| ko04151 | 0.167 | 0.037 | <0.0001 | PI3K-Akt signaling pathway |
| ko04210 | 0.167 | 0.037 | <0.0001 | Apoptosis |
| ko04215 | 0.167 | 0.037 | <0.0001 | Apoptosis - multiple species |
| ko04612 | 0.167 | 0.037 | <0.0001 | Antigen processing and presentation |
| ko04657 | 0.167 | 0.037 | <0.0001 | IL-17 signaling pathway |
| ko04659 | 0.167 | 0.037 | <0.0001 | Th17 cell differentiation |
| ko04914 | 0.167 | 0.037 | <0.0001 | Progesterone-mediated oocyte maturation |
| ko04915 | 0.167 | 0.037 | <0.0001 | Estrogen signaling pathway |
| ko05120 | 0.167 | 0.037 | <0.0001 | Epithelial cell signaling in Helicobacter pylori infection |
| ko05133 | 0.167 | 0.037 | <0.0001 | Pertussis |
| ko05145 | 0.167 | 0.037 | <0.0001 | Toxoplasmosis |
| ko05161 | 0.167 | 0.037 | <0.0001 | Hepatitis B |
| ko05164 | 0.167 | 0.037 | <0.0001 | Influenza A |
| ko05168 | 0.167 | 0.037 | <0.0001 | Herpes simplex infection |
| ko05204 | 0.167 | 0.036 | <0.0001 | Chemical carcinogenesis |
| ko05210 | 0.167 | 0.037 | <0.0001 | Colorectal cancer |
| ko05215 | 0.167 | 0.037 | <0.0001 | Prostate cancer |
| ko05222 | 0.167 | 0.037 | <0.0001 | Small cell lung cancer |
| ko05231 | 0.167 | 0.037 | <0.0001 | Choline metabolism in cancer |
| ko05416 | 0.167 | 0.037 | <0.0001 | Viral myocarditis |
| ko01230 | -0.082 | 0.018 | <0.0001 | Biosynthesis of amino acids |
| ko04922 | -0.215 | 0.047 | <0.0001 | Glucagon signaling pathway |
| ko00960 | 0.101 | 0.022 | <0.0001 | Tropane, piperidine and pyridine alkaloid biosynthesis |
| ko01212 | -0.066 | 0.015 | <0.0001 | Fatty acid metabolism |
| ko00620 | -0.198 | 0.044 | <0.0001 | Pyruvate metabolism |
| ko05200 | 0.123 | 0.028 | <0.0001 | Pathways in cancer |
| ko01055 | 0.101 | 0.023 | <0.0001 | Biosynthesis of vancomycin group antibiotics |
| ko04016 | -0.109 | 0.025 | <0.0001 | MAPK signaling pathway - plant |
| ko01523 | 0.024 | 0.005 | <0.0001 | Antifolate resistance |
| ko00220 | 0.038 | 0.009 | <0.0001 | Arginine biosynthesis |

| | | | | |
|---|---|---|---|---|
| ko00400 | -0.585 | 0.133 | <0.0001 | Phenylalanine, tyrosine and tryptophan biosynthesis |
| ko00750 | 0.051 | 0.012 | <0.0001 | Vitamin B6 metabolism |
| ko04152 | 0.101 | 0.023 | <0.0001 | AMPK signaling pathway |
| ko01100 | -0.016 | 0.004 | <0.0001 | Metabolic pathways |
| ko00785 | 0.045 | 0.010 | <0.0001 | Lipoic acid metabolism |
| ko05134 | 0.104 | 0.025 | <0.0001 | Legionellosis |
| ko01200 | -0.021 | 0.005 | <0.0001 | Carbon metabolism |
| ko00350 | -0.373 | 0.088 | <0.0001 | Tyrosine metabolism |
| ko04112 | 0.085 | 0.020 | <0.0001 | Cell cycle - Caulobacter |
| ko02010 | -0.215 | 0.051 | <0.0001 | ABC transporters |
| ko00520 | -0.306 | 0.074 | <0.0001 | Amino sugar and nucleotide sugar metabolism |
| ko00240 | 0.024 | 0.006 | <0.0001 | Pyrimidine metabolism |
| ko04141 | 0.036 | 0.009 | <0.0001 | Protein processing in endoplasmic reticulum |
| ko03018 | 0.018 | 0.004 | <0.0001 | RNA degradation |
| ko00680 | -0.133 | 0.033 | <0.0001 | Methane metabolism |
| ko00473 | -0.219 | 0.054 | <0.0001 | D-Alanine metabolism |
| ko03060 | 0.078 | 0.019 | <0.0001 | Protein export |
| ko05230 | -0.131 | 0.033 | <0.0001 | Central carbon metabolism in cancer |
| ko00270 | -0.093 | 0.024 | <0.0001 | Cysteine and methionine metabolism |
| ko05010 | 0.144 | 0.037 | <0.0001 | Alzheimer's disease |
| ko04260 | 0.167 | 0.043 | <0.0001 | Cardiac muscle contraction |
| ko04932 | 0.167 | 0.044 | <0.0001 | Non-alcoholic fatty liver disease (NAFLD) |
| ko05012 | 0.167 | 0.044 | <0.0001 | Parkinson's disease |
| ko05211 | -0.031 | 0.008 | <0.0001 | Renal cell carcinoma |
| ko05016 | 0.167 | 0.045 | 0.001 | Huntington's disease |
| ko03070 | 0.131 | 0.035 | 0.001 | Bacterial secretion system |
| ko00561 | -0.154 | 0.042 | 0.001 | Glycerolipid metabolism |
| ko00640 | -0.114 | 0.031 | 0.001 | Propanoate metabolism |
| ko00860 | 0.157 | 0.043 | 0.001 | Porphyrin and chlorophyll metabolism |
| ko02024 | -0.143 | 0.039 | 0.001 | Quorum sensing |
| ko00710 | 0.061 | 0.017 | 0.001 | Carbon fixation in photosynthetic organisms |

| | | | | |
|---|---|---|---|---|
| ko00190 | 0.123 | 0.034 | 0.001 | Oxidative phosphorylation |
| ko01501 | -0.198 | 0.056 | 0.001 | beta-Lactam resistance |
| ko00330 | -0.108 | 0.031 | 0.001 | Arginine and proline metabolism |
| ko00900 | 0.013 | 0.004 | 0.001 | Terpenoid backbone biosynthesis |
| ko01503 | -0.178 | 0.052 | 0.001 | Cationic antimicrobial peptide (CAMP) resistance |
| ko00250 | -0.013 | 0.004 | 0.001 | Alanine, aspartate and glutamate metabolism |
| ko00020 | 0.112 | 0.033 | 0.002 | Citrate cycle (TCA cycle) |
| ko03430 | 0.020 | 0.006 | 0.002 | Mismatch repair |
| ko00760 | -0.251 | 0.077 | 0.003 | Nicotinate and nicotinamide metabolism |
| ko04940 | -0.060 | 0.019 | 0.003 | Type I diabetes mellitus |
| ko04727 | -0.060 | 0.019 | 0.003 | GABAergic synapse |
| ko00650 | -0.108 | 0.034 | 0.003 | Butanoate metabolism |
| ko03420 | 0.016 | 0.005 | 0.003 | Nucleotide excision repair |
| ko00460 | -0.135 | 0.044 | 0.005 | Cyanoamino acid metabolism |
| ko00030 | -0.164 | 0.054 | 0.005 | Pentose phosphate pathway |
| ko00430 | -0.281 | 0.092 | 0.005 | Taurine and hypotaurine metabolism |
| ko00790 | -0.073 | 0.025 | 0.006 | Folate biosynthesis |
| ko03410 | -0.012 | 0.004 | 0.010 | Base excision repair |
| ko00730 | -0.052 | 0.018 | 0.010 | Thiamine metabolism |
| ko01210 | 0.095 | 0.035 | 0.013 | 2-Oxocarboxylic acid metabolism |
| ko00360 | -0.059 | 0.022 | 0.013 | Phenylalanine metabolism |
| ko00040 | -0.199 | 0.075 | 0.015 | Pentose and glucuronate interconversions |
| ko00523 | -0.096 | 0.037 | 0.018 | Polyketide sugar unit biosynthesis |
| ko00983 | -0.098 | 0.038 | 0.018 | Drug metabolism - other enzymes |
| ko00401 | 0.039 | 0.016 | 0.027 | Novobiocin biosynthesis |
| ko00940 | 0.028 | 0.012 | 0.033 | Phenylpropanoid biosynthesis |
| ko01130 | -0.019 | 0.008 | 0.048 | Biosynthesis of antibiotics |

**Table S17 Effect of grouping on functional pathway abundance for male flies for mixed age groups.** Predicted gene function was performed using Piphillin and the KEGG reference database (May 2017 release) using a sequence identity cut-off of 97%. DESeq2 analysis was performed to identify pathways that were differentially abundant between 49-day old single males and males kept in groups of 10 in which focal flies were constantly 1-7 days old. P-values are corrected for multiple testing using the Benjamini-Hochberg method.

| KEGG number | Log2 Fold Change | LFC Standard Error | Adj. P Value | Pathway Function |
|---|---|---|---|---|
| ko00980 | -0.102 | 0.014 | <0.0001 | Metabolism of xenobiotics by cytochrome P450 |
| ko00982 | -0.102 | 0.014 | <0.0001 | Drug metabolism - cytochrome P450 |
| ko01040 | -0.102 | 0.014 | <0.0001 | Biosynthesis of unsaturated fatty acids |
| ko00480 | -0.028 | 0.004 | <0.0001 | Glutathione metabolism |
| ko00908 | 0.075 | 0.011 | <0.0001 | Zeatin biosynthesis |
| ko00910 | -0.133 | 0.020 | <0.0001 | Nitrogen metabolism |
| ko03008 | 0.075 | 0.011 | <0.0001 | Ribosome biogenesis in eukaryotes |
| ko04724 | 0.075 | 0.011 | <0.0001 | Glutamatergic synapse |
| ko05205 | 0.075 | 0.011 | <0.0001 | Proteoglycans in cancer |
| ko01120 | -0.060 | 0.009 | <0.0001 | Microbial metabolism in diverse environments |
| ko04068 | 0.075 | 0.012 | <0.0001 | FoxO signaling pathway |
| ko04211 | 0.075 | 0.012 | <0.0001 | Longevity regulating pathway |
| ko05014 | 0.075 | 0.012 | <0.0001 | Amyotrophic lateral sclerosis (ALS) |
| ko00051 | -0.489 | 0.077 | <0.0001 | Fructose and mannose metabolism |
| ko00521 | -0.190 | 0.030 | <0.0001 | Streptomycin biosynthesis |
| ko02020 | -0.151 | 0.024 | <0.0001 | Two-component system |
| ko00660 | 0.075 | 0.012 | <0.0001 | C5-Branched dibasic acid metabolism |
| ko04626 | 0.075 | 0.012 | <0.0001 | Plant-pathogen interaction |
| ko00260 | -0.055 | 0.009 | <0.0001 | Glycine, serine and threonine metabolism |
| ko00061 | -0.088 | 0.014 | <0.0001 | Fatty acid biosynthesis |
| ko04213 | 0.075 | 0.012 | <0.0001 | Longevity regulating pathway - multiple species |

| | | | | |
|---|---|---|---|---|
| ko00310 | 0.113 | 0.018 | <0.0001 | Lysine degradation |
| ko04122 | -0.078 | 0.012 | <0.0001 | Sulfur relay system |
| ko05111 | -0.418 | 0.067 | <0.0001 | Biofilm formation - Vibrio cholerae |
| ko04214 | 0.113 | 0.018 | <0.0001 | Apoptosis - fly |
| ko05152 | 0.075 | 0.012 | <0.0001 | Tuberculosis |
| ko04066 | 0.056 | 0.009 | <0.0001 | HIF-1 signaling pathway |
| ko04212 | 0.065 | 0.011 | <0.0001 | Longevity regulating pathway - worm |
| ko02026 | -0.370 | 0.061 | <0.0001 | Biofilm formation - Escherichia coli |
| ko00920 | -0.346 | 0.057 | <0.0001 | Sulfur metabolism |
| ko00010 | -0.187 | 0.031 | <0.0001 | Glycolysis / Gluconeogenesis |
| ko00770 | -0.164 | 0.028 | <0.0001 | Pantothenate and CoA biosynthesis |
| ko00400 | -0.787 | 0.133 | <0.0001 | Phenylalanine, tyrosine and tryptophan biosynthesis |
| ko00130 | 0.086 | 0.015 | <0.0001 | Ubiquinone and other terpenoid-quinone biosynthesis |
| ko01100 | -0.022 | 0.004 | <0.0001 | Metabolic pathways |
| ko00280 | 0.063 | 0.011 | <0.0001 | Valine, leucine and isoleucine degradation |
| ko01230 | -0.106 | 0.018 | <0.0001 | Biosynthesis of amino acids |
| ko00740 | 0.056 | 0.010 | <0.0001 | Riboflavin metabolism |
| ko01524 | 0.136 | 0.023 | <0.0001 | Platinum drug resistance |
| ko04922 | -0.275 | 0.047 | <0.0001 | Glucagon signaling pathway |
| ko04070 | 0.115 | 0.020 | <0.0001 | Phosphatidylinositol signaling system |
| ko04621 | 0.038 | 0.007 | <0.0001 | NOD-like receptor signaling pathway |
| ko00620 | -0.253 | 0.044 | <0.0001 | Pyruvate metabolism |
| ko01212 | -0.084 | 0.015 | <0.0001 | Fatty acid metabolism |
| ko03030 | 0.051 | 0.009 | <0.0001 | DNA replication |
| ko00630 | 0.067 | 0.012 | <0.0001 | Glyoxylate and dicarboxylate metabolism |
| ko04146 | 0.090 | 0.016 | <0.0001 | Peroxisome |
| ko00564 | 0.047 | 0.008 | <0.0001 | Glycerophospholipid metabolism |
| ko00670 | 0.043 | 0.008 | <0.0001 | One carbon pool by folate |
| ko00195 | 0.088 | 0.016 | <0.0001 | Photosynthesis |
| ko00950 | 0.203 | 0.037 | <0.0001 | Isoquinoline alkaloid biosynthesis |

| | | | | |
|---|---|---|---|---|
| ko01200 | -0.028 | 0.005 | <0.0001 | Carbon metabolism |
| ko03440 | 0.068 | 0.012 | <0.0001 | Homologous recombination |
| ko04013 | 0.203 | 0.037 | <0.0001 | MAPK signaling pathway - fly |
| ko04016 | -0.137 | 0.025 | <0.0001 | MAPK signaling pathway - plant |
| ko04072 | 0.203 | 0.037 | <0.0001 | Phospholipase D signaling pathway |
| ko04115 | 0.203 | 0.037 | <0.0001 | p53 signaling pathway |
| ko04151 | 0.203 | 0.037 | <0.0001 | PI3K-Akt signaling pathway |
| ko04210 | 0.203 | 0.037 | <0.0001 | Apoptosis |
| ko04215 | 0.203 | 0.037 | <0.0001 | Apoptosis - multiple species |
| ko04612 | 0.203 | 0.037 | <0.0001 | Antigen processing and presentation |
| ko04657 | 0.203 | 0.037 | <0.0001 | IL-17 signaling pathway |
| ko04659 | 0.203 | 0.037 | <0.0001 | Th17 cell differentiation |
| ko04914 | 0.203 | 0.037 | <0.0001 | Progesterone-mediated oocyte maturation |
| ko04915 | 0.203 | 0.037 | <0.0001 | Estrogen signaling pathway |
| ko05120 | 0.202 | 0.037 | <0.0001 | Epithelial cell signaling in Helicobacter pylori infection |
| ko05133 | 0.202 | 0.037 | <0.0001 | Pertussis |
| ko05145 | 0.203 | 0.037 | <0.0001 | Toxoplasmosis |
| ko05161 | 0.203 | 0.037 | <0.0001 | Hepatitis B |
| ko05164 | 0.203 | 0.037 | <0.0001 | Influenza A |
| ko05168 | 0.203 | 0.037 | <0.0001 | Herpes simplex infection |
| ko05204 | 0.202 | 0.036 | <0.0001 | Chemical carcinogenesis |
| ko05210 | 0.203 | 0.037 | <0.0001 | Colorectal cancer |
| ko05215 | 0.203 | 0.037 | <0.0001 | Prostate cancer |
| ko05222 | 0.203 | 0.037 | <0.0001 | Small cell lung cancer |
| ko05231 | 0.203 | 0.037 | <0.0001 | Choline metabolism in cancer |
| ko05416 | 0.203 | 0.037 | <0.0001 | Viral myocarditis |
| ko05418 | 0.064 | 0.012 | <0.0001 | Fluid shear stress and atherosclerosis |
| ko03010 | 0.047 | 0.009 | <0.0001 | Ribosome |
| ko00350 | -0.483 | 0.088 | <0.0001 | Tyrosine metabolism |
| ko00960 | 0.120 | 0.022 | <0.0001 | Tropane, piperidine and pyridine alkaloid biosynthesis |

| | | | | |
|---|---|---|---|---|
| ko05200 | 0.149 | 0.028 | <0.0001 | Pathways in cancer |
| ko02010 | -0.275 | 0.051 | <0.0001 | ABC transporters |
| ko00520 | -0.394 | 0.074 | <0.0001 | Amino sugar and nucleotide sugar metabolism |
| ko01055 | 0.120 | 0.023 | <0.0001 | Biosynthesis of vancomycin group antibiotics |
| ko04152 | 0.120 | 0.023 | <0.0001 | AMPK signaling pathway |
| ko00220 | 0.045 | 0.009 | <0.0001 | Arginine biosynthesis |
| ko00471 | 0.016 | 0.003 | <0.0001 | D-Glutamine and D-glutamate metabolism |
| ko05134 | 0.126 | 0.025 | <0.0001 | Legionellosis |
| ko00750 | 0.059 | 0.012 | <0.0001 | Vitamin B6 metabolism |
| ko00473 | -0.278 | 0.054 | <0.0001 | D-Alanine metabolism |
| ko00680 | -0.170 | 0.033 | <0.0001 | Methane metabolism |
| ko04112 | 0.103 | 0.020 | <0.0001 | Cell cycle - Caulobacter |
| ko00785 | 0.053 | 0.010 | <0.0001 | Lipoic acid metabolism |
| ko00270 | -0.120 | 0.024 | <0.0001 | Cysteine and methionine metabolism |
| ko05230 | -0.166 | 0.033 | <0.0001 | Central carbon metabolism in cancer |
| ko01523 | 0.026 | 0.005 | <0.0001 | Antifolate resistance |
| ko03060 | 0.094 | 0.019 | <0.0001 | Protein export |
| ko00240 | 0.028 | 0.006 | <0.0001 | Pyrimidine metabolism |
| ko05010 | 0.176 | 0.037 | <0.0001 | Alzheimer's disease |
| ko00250 | -0.018 | 0.004 | <0.0001 | Alanine, aspartate and glutamate metabolism |
| ko05211 | -0.037 | 0.008 | <0.0001 | Renal cell carcinoma |
| ko04141 | 0.040 | 0.009 | <0.0001 | Protein processing in endoplasmic reticulum |
| ko04260 | 0.203 | 0.043 | <0.0001 | Cardiac muscle contraction |
| ko00561 | -0.195 | 0.042 | <0.0001 | Glycerolipid metabolism |
| ko00640 | -0.144 | 0.031 | <0.0001 | Propanoate metabolism |
| ko02024 | -0.182 | 0.039 | <0.0001 | Quorum sensing |
| ko04932 | 0.203 | 0.044 | <0.0001 | Non-alcoholic fatty liver disease (NAFLD) |
| ko05012 | 0.203 | 0.044 | <0.0001 | Parkinson's disease |
| ko03018 | 0.020 | 0.004 | <0.0001 | RNA degradation |
| ko03070 | 0.160 | 0.035 | <0.0001 | Bacterial secretion system |
| ko05016 | 0.203 | 0.045 | <0.0001 | Huntington's disease |
| ko01501 | -0.252 | 0.056 | <0.0001 | beta-Lactam resistance |

| | | | | |
|---|---|---|---|---|
| ko00860 | 0.191 | 0.043 | <0.0001 | Porphyrin and chlorophyll metabolism |
| ko00330 | -0.137 | 0.031 | <0.0001 | Arginine and proline metabolism |
| ko00710 | 0.073 | 0.017 | <0.0001 | Carbon fixation in photosynthetic organisms |
| ko00190 | 0.150 | 0.034 | <0.0001 | Oxidative phosphorylation |
| ko01503 | -0.226 | 0.052 | <0.0001 | Cationic antimicrobial peptide (CAMP) resistance |
| ko00760 | -0.319 | 0.077 | <0.0001 | Nicotinate and nicotinamide metabolism |
| ko00020 | 0.136 | 0.033 | <0.0001 | Citrate cycle (TCA cycle) |
| ko00650 | -0.137 | 0.034 | <0.0001 | Butanoate metabolism |
| ko04940 | -0.075 | 0.019 | <0.0001 | Type I diabetes mellitus |
| ko04727 | -0.075 | 0.019 | <0.0001 | GABAergic synapse |
| ko03410 | -0.017 | 0.004 | <0.0001 | Base excision repair |
| ko00430 | -0.357 | 0.092 | <0.0001 | Taurine and hypotaurine metabolism |
| ko00030 | -0.207 | 0.054 | <0.0001 | Pentose phosphate pathway |
| ko00790 | -0.096 | 0.025 | <0.0001 | Folate biosynthesis |
| ko00460 | -0.170 | 0.044 | <0.0001 | Cyanoamino acid metabolism |
| ko03430 | 0.023 | 0.006 | <0.0001 | Mismatch repair |
| ko00900 | 0.013 | 0.004 | 0.001 | Terpenoid backbone biosynthesis |
| ko00730 | -0.066 | 0.018 | 0.001 | Thiamine metabolism |
| ko00360 | -0.075 | 0.022 | 0.001 | Phenylalanine metabolism |
| ko03420 | 0.017 | 0.005 | 0.001 | Nucleotide excision repair |
| ko00523 | -0.125 | 0.037 | 0.001 | Polyketide sugar unit biosynthesis |
| ko00040 | -0.251 | 0.075 | 0.001 | Pentose and glucuronate interconversions |
| ko01210 | 0.115 | 0.035 | 0.002 | 2-Oxocarboxylic acid metabolism |
| ko00983 | -0.124 | 0.038 | 0.002 | Drug metabolism - other enzymes |
| ko01130 | -0.026 | 0.008 | 0.004 | Biosynthesis of antibiotics |
| ko00401 | 0.044 | 0.016 | 0.009 | Novobiocin biosynthesis |
| ko00940 | 0.032 | 0.012 | 0.012 | Phenylpropanoid biosynthesis |
| ko00450 | -0.031 | 0.012 | 0.015 | Selenocompound metabolism |
| ko00562 | -0.015 | 0.006 | 0.024 | Inositol phosphate metabolism |
| ko00290 | -2.181 | 0.929 | 0.032 | Valine, leucine and isoleucine biosynthesis |
| ko00332 | -2.418 | 1.038 | 0.034 | Carbapenem biosynthesis |

| | | | | |
|---|---|---|---|---|
| ko00340 | -2.417 | 1.040 | 0.034 | Histidine metabolism |
| ko02025 | -2.420 | 1.050 | 0.035 | Biofilm formation - Pseudomonas aeruginosa |
| ko00405 | -2.420 | 1.055 | 0.036 | Phenazine biosynthesis |
| ko00623 | -2.052 | 0.904 | 0.038 | Toluene degradation |
| ko00603 | -2.186 | 0.969 | 0.040 | Glycosphingolipid biosynthesis - globo and isoglobo series |
| ko00311 | -2.189 | 0.987 | 0.043 | Penicillin and cephalosporin biosynthesis |
| ko00500 | -2.303 | 1.040 | 0.043 | Starch and sucrose metabolism |
| ko00052 | -2.225 | 1.007 | 0.044 | Galactose metabolism |
| ko00440 | -2.251 | 1.022 | 0.044 | Phosphonate and phosphinate metabolism |
| ko00622 | -2.058 | 0.937 | 0.045 | Xylene degradation |
| ko00970 | 0.013 | 0.006 | 0.045 | Aminoacyl-tRNA biosynthesis |
| ko01054 | -2.138 | 0.977 | 0.045 | Nonribosomal peptide structures |
| ko01220 | -2.137 | 0.978 | 0.045 | Degradation of aromatic compounds |
| ko02060 | -2.385 | 1.099 | 0.047 | Phosphotransferase system (PTS) |
| ko00524 | -2.059 | 0.951 | 0.047 | Neomycin, kanamycin and gentamicin biosynthesis |
| ko04918 | -2.158 | 1.008 | 0.049 | Thyroid hormone synthesis |
| ko00830 | -2.140 | 1.002 | 0.050 | Retinol metabolism |

-infected food eaten by males and females kept singly and in same-sex pairs.
